# High-throughput evolution of near-infrared oxytocin nanosensors enables oxytocin imaging in mice and prairie voles

**DOI:** 10.1101/2024.05.10.593556

**Authors:** Jaquesta A.M. Adams, Natsumi Komatsu, Nicole Navarro, Alexis M. Black, Esther Leem, Xiaoqi Sun, Jiaxuan Zhao, Octavio I. Arias-Soto, Annaliese K. Beery, Markita P. Landry

## Abstract

Oxytocin is a neuropeptide involved in regulating social and emotional behavior. Current techniques for oxytocin imaging are generally limited in spatial and temporal resolution, real-time imaging capacity, selectivity for oxytocin over vasopressin, and application in young and non-model organisms. To address these issues, we developed a method to evolve purely synthetic molecular recognition for oxytocin on the surface of near-infrared fluorescent single-walled carbon nanotubes (SWCNT) using single-stranded DNA (ssDNA). The best-performing nanosensor nIROT-SELEC reversibly undergoes up to a 172% fluorescence increase in response to oxytocin with micromolar dissociation, nanomolar limit of detection, and and high selectivity over oxytocin analogs, receptor agonists and antagonists, and co-released neurochemicals. We next demonstrated the versatility of nIROT-SELEC by performing live imaging of synaptic evoked oxytocin released in acute brain slices of mice and prairie voles. Our method for high throughput evolution of neuropeptide nanosensors holds promise to enable synaptic scale visualization of neuropeptide signaling in the brain cross different species and developmental stages, to advance the study of neurochemical signaling for its role in both health and disease.

## Introduction

Neuropeptides have been hypothesized to play a central role over a broad range of neurotypical conditions with aberrations in their signaling leading to developmental and psychiatric disorders. However, imaging neuropeptides has remained challenging, especially in non-model organisms that uniquely exhibit behaviors relevant to neuropeptide signaling. One such neuropeptide, oxytocin, is a nonapeptide known for its central role in regulating social and emotional behaviors ^1,2^. Historically, oxytocin was first recognized for its influence in reproductive actions such as lactation and uterine contraction, but has since been implicated in behaviors such as recognition, pair and maternal bonding, non-reproductive social interactions, and anxiolysis ^3–7^. Consequently, dysfunction in oxytocin signaling has been implicated in the pathogenesis of neurological disorders related to social impairment such as schizophrenia, mood disorders, anxiety, and autism spectrum disorder^8,9^. Understanding oxytocin signaling is therefore of paramount interest in social neuroscience^3,4,11,12^. For instance, prairie voles exhibit higher levels of oxytocin receptor expression in the nucleus accumbens compared to mice, a factor considered to influence their unique social behaviors, including social monogamy^13^. However, understanding of how oxytocin supports variation in complex social behaviors across species, as well as its role in both health and disease, remains incomplete owing to a lack of versatile methods to directly probe oxytocin signaling at the spatiotemporal resolution requisite to elucidate oxytocinergic communication.

Microdialysis coupled with an assay has been widely used to selectively measure neuropeptides such as oxytocin, yet sampling typically occurs over minutes, wherein sensitive samples may degrade and transient changes will not be captured^14,15^. By leveraging the redox active tyrosine, electrochemical methods can achieve rapid detection of oxytocin, albeit with lower sensitivity relative to immunoassay-coupled microdialysis^16^. Recently, genetically encoded oxytocin sensing platforms have been reported for oxytocin^17,18^. However, the timescale required for uniform expression of genetically encoded sensors may present a challenge in studying oxytocin release in young organisms, which is a key time point for the study of social development and disorders thereof. This leaves a need for real-time oxytocin imaging probes that can distinguish oxytocin from vasopressin and can be used irrespective of an organism’s species and age, and as of now, no real-time oxytocin imaging in non-model animals has been demonstrated. This critical gap motivates the development of synthetic optical probes that can be applied across species and developmental stages to add to the oxytocin interrogative toolkit.

To this end, we have developed a synthetic molecular evolution method, systematic evolution of ssDNA ligands by exponential enrichment following adsorption to carbon nanotubes (SELEC) ^19^, a generic method that can be used to evolve synthetic molecular recognition for any neuropeptide target. Here, we demonstrate the utility of SELEC to evolve a nanosensor for synaptic-scale oxytocin imaging. The probe leverages the inherent tissue-transparent and infinitely photostable fluorescence of single-walled carbon nanotubes (SWCNT) and molecular recognition of the DNA-SWCNT construct. While peptide-based nansensors for oxytocin have recently been developed^24^, our SELEC based approach offers unique advantages of facile synthesis, superior sensitivity, and a purely synthetic platform that can be generically used to develop nanosensors for any neuropeptide. Here, we present nIROT-SELEC, a nanosensor which responds reversibly and selectively to oxytocin with a change in near-infrared fluorescence, ΔF/F_0_, of up to 172% and a K_d_ of 4.93±3.19 μM (R_2_ > 0.99) *in vitro*. We show that nIROT-SELEC can be used to image electrically stimulated oxytocin release in acute brain tissue slices in mice and in prairie voles, suggesting that nIROT-SELEC and broadly the SELEC method can serve as a valuable approach to studying neuropeptide signaling.

## Results

### Validation of SELEC as a method to evolve synthetic molecular recognition for neuropeptides, and applied to oxytocin

We implemented SELEC to evolve ssDNA-SWCNT molecular recognition for oxytocin by designing our initial ssDNA library to be composed of greater than 10^10^ random 18-mer ssDNA sequences flanked by two C_6_-mers and two 18-mer primer regions for polymerase chain reaction (PCR) amplification (Fig. 1). The PCR primers enabled enzymatic amplification and enrichment of high-affinity ssDNA. The purpose of the C_6_ sequences was twofold: (1) the C_6_ sequence has a high affinity for the carbon nanotube surface thus pinning the ssDNA sequences to the SWCNT and (2) the 6-mer sequences separate the random binding region of the ssDNA from the primer regions ^25^. Briefly, 10 µg of SWCNT was suspended using 100 nmol of the library of unique ssDNA sequences (200 µL, 0.5 mM), 100 nmol of oxytocin (100 µL, 1 mM), and 200 µL 20 mM acetate buffer to maintain a pH of 5 for oxytocin structural stability. An excess of ssDNA was used to promote competitive complexation of ssDNA to SWCNT in a manner that promotes oxytocin selectivity in the experimental library. Following competitive complexation, unbound ssDNA were removed by spin filtration. Bound ssDNA were subsequently desorbed, amplified by PCR, and isolated prior to use in subsequent rounds of SELEC. We determined that six rounds of SELEC is a sufficient threshold to observe neurochemical molecular recognition in the evolved library by principal component analysis of sequence similarity in the experimental versus control SELEC groups, which showed divergence by six rounds of evolution (Fig. 2a and b). ssDNA sequences from rounds 3-6 of SELEC were prepared and submitted for deep sequencing with an Illumina HiSeq 4000. A control sample was generated similarly but without an analyte as described previously ^19^. Control SELEC was used to identify ssDNA that only have a high affinity for the SWCNT surface, not the target analyte, or are subject to PCR biases ^26^. Sequencing analysis of the top oxytocin SELEC nanosensor candidates utilized only the 18-mer random region, excluding the polycytosine and PCR primer regions. *In vitro* and *ex vivo* characterization of the top candidates utilized the 18-mer random region and the C_6_ regions to increase the density of molecular recognition sites on the SWCNT surface. Previous work demonstrated that nanosensors synthesized without the primer regions have increased sensitivity toward the target analyte.^19^

**Figure 1.**
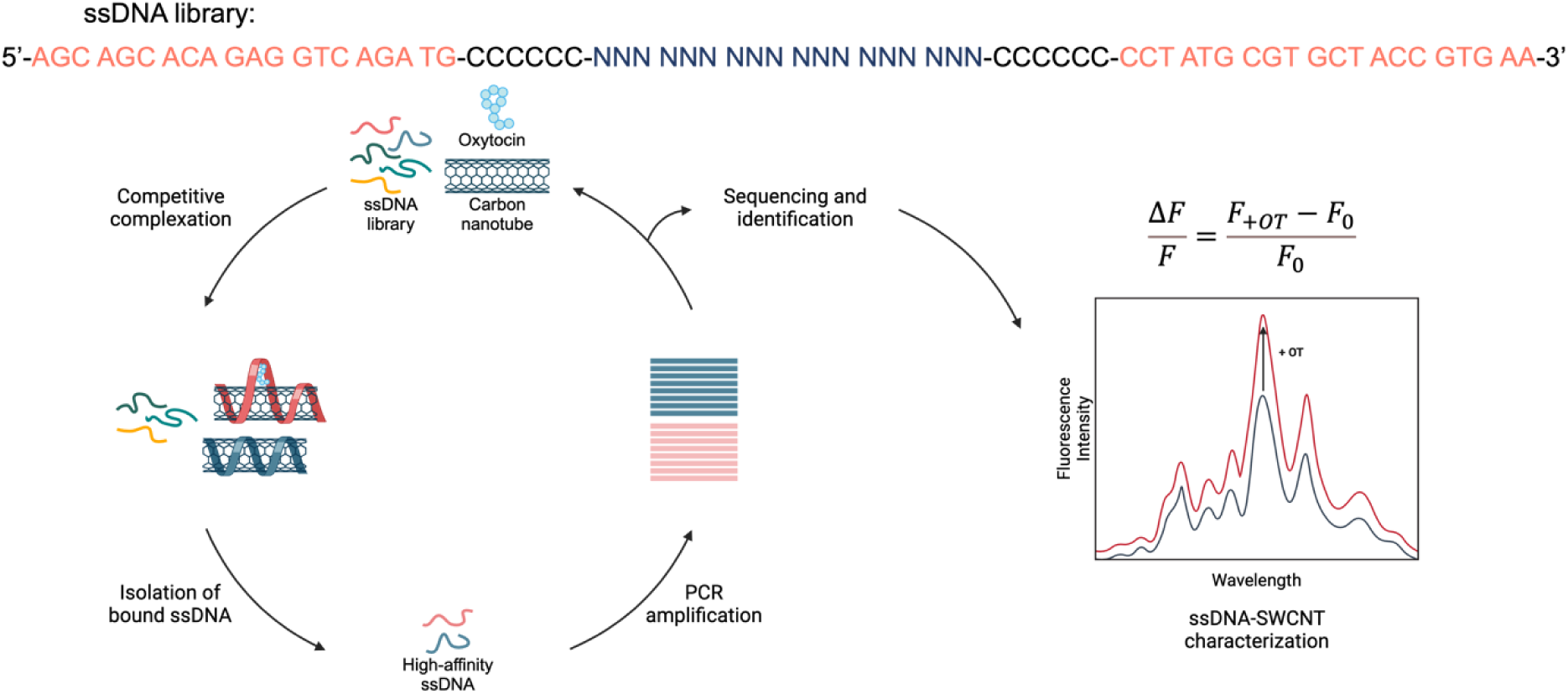
Schematic illustration of the SELEC nanosensor discovery process. In SELEC, SWCNTs are incubated with a ≥10-fold mass excess of ssDNA and 100 μM oxytocin (experimental library) or ssDNA only (control library). These mixtures are tip sonicated to generate a complex of either ssDNA-SWCNT bound to oxytocin (red) or ssDNA-SWCNT alone (blue). Unbound ssDNA are removed by spin-filtration and bound ssDNA are separated from SWCNT by thermal desorption to isolate high-affinity ssDNA. High-affinity ssDNA are then amplified by PCR and fed into the next selection round. Following six rounds, ssDNA selective for oxytocin when bound to SWCNT were identified by high-throughput sequencing and characterized by nIR spectroscopy. Created with BioRender.com.

**Figure 2.**
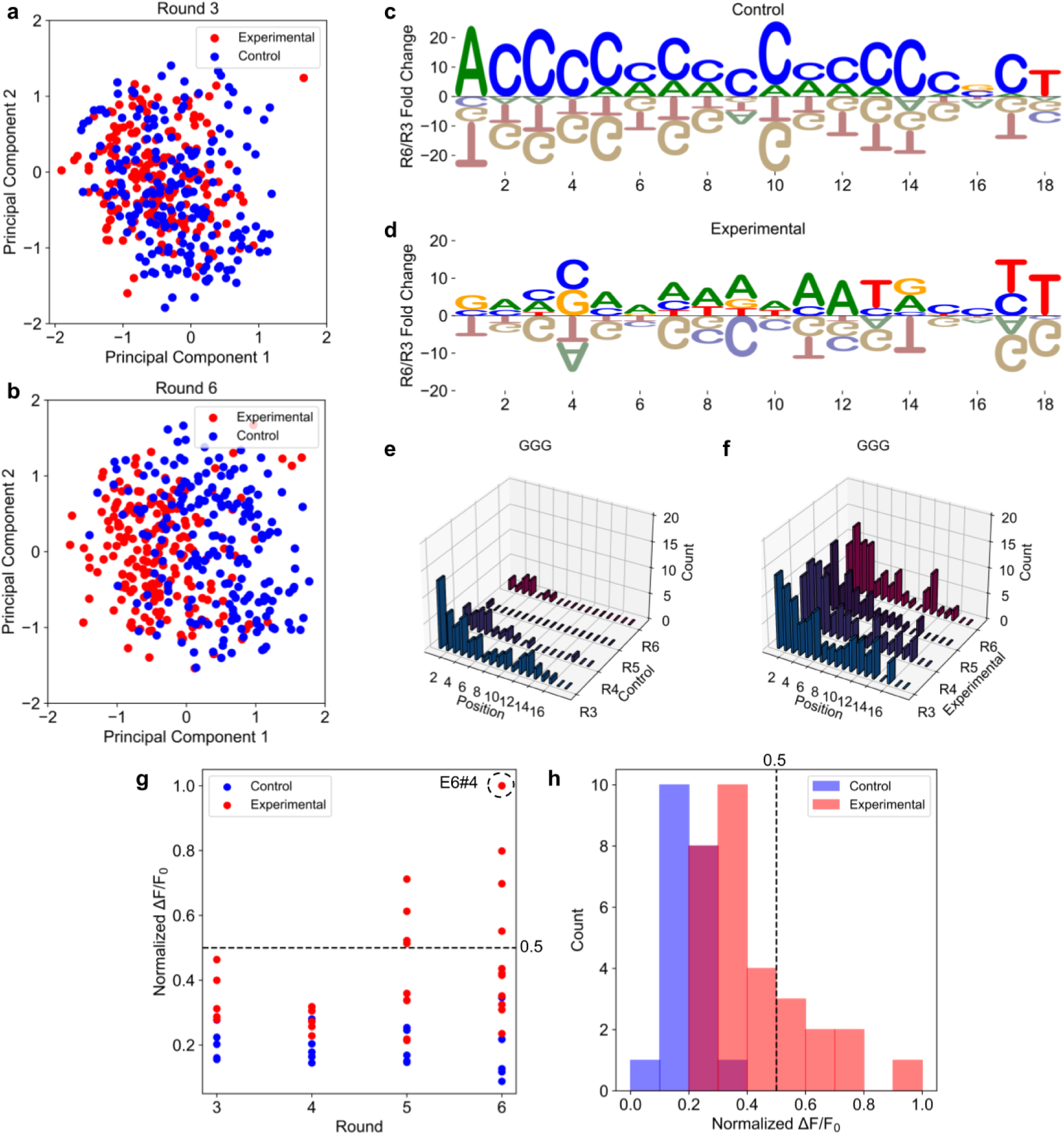
SELEC evolves ssDNA-SWCNT nanosensors for molecular recognition of oxytocin. **(a)** and **(b)** PCA plot for experimental (red) and control (blue) SELEC library sequences. Principal component analysis was performed for the top 200 ranked ssDNA sequences from rounds 3 to 6 of the experimental and control SELEC libraries. Principal component 1 is plotted against principal component 2. **(c)** and **(d)** Fold change for each nucleotide at each position in the experimental library and control library from round 3 to round 6. **(e)** and **(f)** Trimer frequency of GGG at each nucleotide position for the experimental (left) and control (right) libraries across rounds 3 to 6. **(g)** Normalized ΔF/F_0_ upon 100 μM oxytocin addition to ssDNA-SWCNT constructs prepared from the top sequences across experimental and control rounds. Each point represents the average normalized ΔF/F_0_ of n = 3 measurements. The most sensitive nanosensor, E6#4, is circled. Data is visualized as a histogram in **(h).**

We identified the top 200 sequences from SELEC rounds 3 and 6 and analyzed them using principal component analysis (PCA) (Fig. 2a and b). Divergence of the experimental and control libraries was not observed in round 3 but was observed in round 6. This observation, attributable to the evolution of distinct nucleotide compositions, is likely resulting from the presence of oxytocin in SELEC during SELEC evolution, indicating the successful evolution of sequences for oxytocin molecular recognition. To further support this hypothesis, we calculated nucleotide fold change across the positions of the 18-mer random region from round 3 to round 6 in the experimental and control libraries (Fig. 2c and d). Key differences in nucleotide identity as a function of position along the ssDNA strand were identified between nucleotide enrichment in the experimental and control libraries. The control library demonstrated enrichment of cytosine at nearly every position except the first and final positions, as expected from prior literature demonstrating that cytosine is a tight-binding nucleotide for the SWCNT surface ^25^. However, in the experimental library, adenine was the most-enriched nucleotide overall, demonstrating preeminent fold change at positions 2 and 5-12. Furthermore, the experimental library generally demonstrated heterogeneity in nucleotide fold change across positions relative to the control library. Interestingly, both the experimental library and control library were most likely to contain adenine at position 1 and guanine at position 18 (Fig. S1), suggesting these trends at the outermost positions of the random region evolved to facilitate binding of the sequence to SWCNT rather than oxytocin.

We also analyzed the positional frequency of trimer appearance in the top 200 sequences of the experimental and control libraries. GGG was the most enriched trimer in the experimental library relative to the control library. GGG initially appeared in the top 20 trimers of the round 3 control library but declined in prevalence in the rounds that followed (Fig. 2e and f). Similarly, AGG and GGC were enriched across the experimental library and depleted in the control library, implicating these trimers in the evolution of oxytocin molecular recognition during SELEC (Fig. S2). Each of these trimers also displayed positional preference for the 5’ end of the random recognition sequence.

GTG was the most abundant trimer across experimental rounds and the third-most abundant trimer across control rounds (Fig. S3). GTG also displayed positional preference with a strong enrichment on the 3’ end of the random recognition sequence. ACG and GCA were the only other trimers to appear in the top ten most abundant trimers in both the experimental and control libraries with GCA appearing at position 10 in both cases. The presence of these trimers in both the experimental and control libraries suggests that, while they may have a role in oxytocin molecular recognition, they may also be influential in SWCNT binding. The top 10 trimers present across the experimental and control SELEC libraries, respectively, are presented in Table S1 and Table S2 in order of descending frequency.

Following nucleotide composition analysis of the SELEC libraries, we evaluated the sensitivity of ssDNA-SWCNT constructs prepared using the top enriched experimental and control library sequences per round. Sequences are identified as EN#M, where E represents experimental, N represents the SELEC round number (3-6), and M represents the sequence’s frequency ranking within said round (1-10). Correspondingly, control sequences are referred to as CN#M. ssDNA-SWCNT nanosensors were prepared from the top 5 sequences of control SELEC rounds 3-6 and experimental SELEC rounds 3 and 4. The top ten sequences were selected for nanosensor synthesis from rounds 5 and 6 of experimental SELEC as sequences from these rounds were expected to have higher affinity for oxytocin than those from rounds 3 and 4 (Table S3).

ssDNA-SWCNT nanosensors were prepared by mixing ssDNA (100 μM, 1 mM in H_2_O) with SWCNT (900 µL, 0.22 mg/mL in 1X PBS) and noncovalently adsorbing ssDNA to the SWCNT surface by probe-tip sonication. All ssDNA sequences successfully suspended SWCNT. The candidate nanosensors were then diluted to 5 mg/L in PBS before fluorescence spectra were acquired (Fig. S4). Nanosensor optical response is reported as ΔF/F_0_ = (F_t_-F_0_)/ F_0_, where F_0_ is the baseline nanosensor fluorescence and F_t_ is the peak fluorescence after analyte incubation at timepoint *t*. Nanosensor response can be wavelength-dependent. Correspondingly, we observe the highest sensitivity to oxytocin at the peak corresponding to the 1195 nm wavelength and (8,6) chirality SWCNT, likely due to the 721 nm excitation wavelength used to acquire our spectra which excites the (8,6) chirality on-resonance ^27,28^. We therefore use this peak for all peak ΔF/F_0_ calculations.

The normalized fluorescence response of ssDNA-SWCNT nanosensors to 100 µM oxytocin was calculated for the selected top ssDNA sequences from the control and experimental libraries as a function of SELEC round (Fig. 2g). Fluorescence was normalized to the top responder in the experimental SELEC library, E6#4-SWCNT. Nanosensors prepared with control library sequences responded to oxytocin with a mean ΔF/F_0_ = 0.19 ± 0.06, while those prepared with experimental library sequences responded to oxytocin with a mean ΔF/F_0_ = 0.41 ± 0.19. The higher responsivity of nanosensors prepared with sequences from the experimental library compared to those from the control library indicates that these former sequences were successfully evolved via SELEC to bind oxytocin. Furthermore, all ssDNA-SWCNT constructs with a ΔF/F_0_ > 0.5 were identified through experimental SELEC rounds 5 and 6, with E6#4-SWCNT demonstrating the largest optical response to oxytocin [ΔF/F_0_ = 1.05 ± 0.04 (mean ± SD); n = 3] (Fig. 2h). The increased appearance of high-responding sequences in experimental SELEC, particularly rounds 5 and 6, also supports that oxytocin sensitivity evolved as a function of SELEC round, as expected. As the most oxytocin-responsive nanosensor candidate, E6#4-SWCNT, was selected for further characterization as nIROT-SELEC (**n**ear-**I**nfra**R**ed **O**xy**T**ocin nanosensor identified by **SELEC**). The sequence and fluorescence intensity spectrum for nIROT-SELEC in response to oxytocin is shown in Fig. 3a.

**Figure 3.**
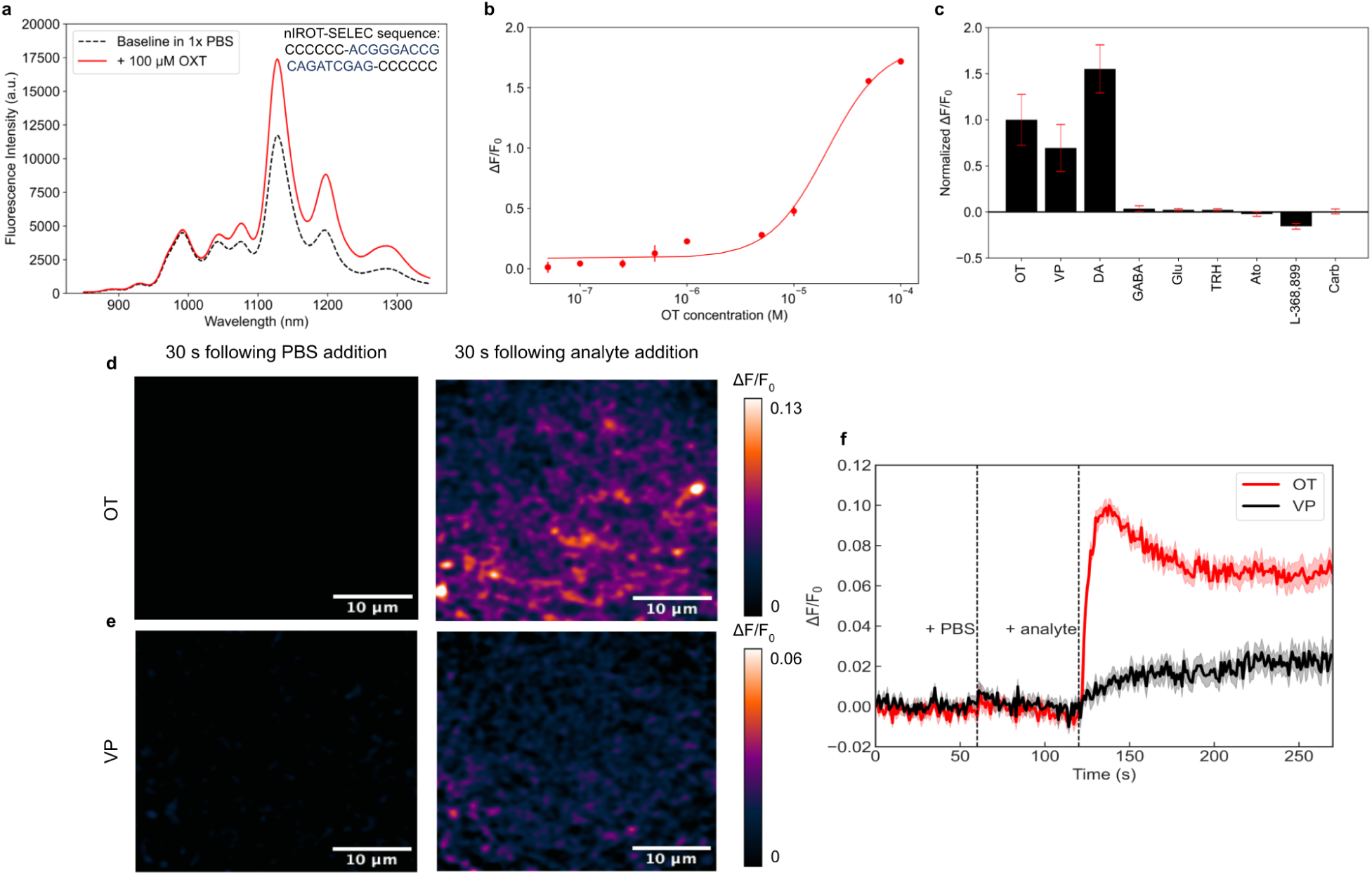
nIROT-SELEC demonstrates oxytocin sensitivity and selectivity. **(a)** A single fluorescence intensity spectra for nIROT-SELEC before (black, dotted) and after (red) addition of 100 μM oxytocin. **(b)** Fluorescence modulation of nIROT-SELEC in response to oxytocin concentrations from 50 nM to 100 μM in PBS. Each point represents n=3 measurements. **(c)** Response of nIROT-SELEC to 50 µM of oxytocin (OT) (n=15), vasopressin (VP) (n=6), dopamine (DA) (n = 3), y-aminobutyric acid (GABA) (n = 3), glutamate (Glu) (n = 3), thyrotropin hormone (TRH) (n = 3), atosiban (Ato) (n = 3), L-368,899 (n = 3), and carbetocin (Carb) (n = 3). Error bars are standard deviation. **(d)** nIR fluorescence images of immobilized nIROT-SELEC in the same field of view (left) 30 seconds following PBS addition and (right) 30 seconds following oxytocin addition. **(e)** nIR fluorescence images of immobilized nIROT-SELEC in the same field of view (left) 30 seconds following PBS addition and (right) 30 seconds following vasopressin addition. Scale bars represent 10 µm. **(f)** Time-course integrated ΔF/F_0_ of immobilized nIROT-SELEC to 100 µM oxytocin (red line) and 100 µM vasopressin (black line). The line is the average of 10 ROIs. The shaded area is standard deviation.

### Characterization of nIROT-SELEC nanosensor selectivity over vasopressin

We next characterized our nIROT-SELEC nanosensor for possible in-brain oxytocin imaging via *in vitro* characterization by spectroscopically measuring its response towards varying concentrations of oxytocin (Fig. 3b). nIROT-SELEC responded to 76-minute-long incubation with oxytocin in a concentration-dependent manner with nanomolar sensitivity. We next calculated nanosensor kinetic parameters by fitting nIROT-SELEC fluorescence response versus oxytocin concentration to a cooperative binding model ^29^. The equilibrium dissociation constant (K_d_) was determined to be 4.93 ± 3.19 μM. To characterize nanosensor selectivity, nIROT-SELEC fluorescence modulation was evaluated for response to 50 µM concentrations of a panel of neurochemicals after 20 minutes, including vasopressin, dopamine, ɣ-aminobutyric acid (GABA), glutamate, and thyrotropin-releasing hormone (TRH) (Fig. 3c). nIROT-SELEC is selective against TRH [ΔF/F_0_ = 0.024 ± 0.013 (mean ± SD); n = 3], which is co-released in the hypothalamus alongside oxytocin.^29,30^ As expected, nIROT-SELEC responds significantly to dopamine relative to oxytocin [ΔF/F_0_ = 1.55 ± 0.26 (mean ± SD); n = 3]. Nearly all ssDNA-SWCNT constructs demonstrate optical sensitivity to dopamine, necessitating that nIROT-SELEC is exclusively applied in nondopaminergic brain regions or that dopamine-suppressing pharmacology is implemented when probing dopaminergic regions. When using fluorescence spectroscopy, nIROT-SELEC is moderately sensitive to the oxytocin analog vasopressin relative to oxytocin [ΔF/F_0_ = 0.69 ± 0.25 (mean ± SD); n=6]. To further explore the selectivity of nIROT-SELEC for oxytocin over vasopressin in the imaging form factor in which its use is intended, we surface-immobilized nIROT-SELEC on a glass slide and imaged these surface-immobilized nanosensors prior to and following analyte addition (Fig. 3d, e, and f). Nanosensors were immobilized on MatTek glass-bottom microwell dishes (35 mm petri dish with 10 mm microwell). PBS was added after 60 s, and either oxytocin or vasopressin was added after 120 s. Following PBS addition, the average ΔF/F_0_ of 20 nanosensor regions of interest (ROIs) in a single field of view was −0.002 ± 0.003 (mean ± SD). When washed with oxytocin, the average ΔF/F_0_ of 10 ROIs in a single field of view was 0.091 ± 0.004 (mean ± SD) after 30 seconds. When washed with vasopressin, the average ΔF/F_0_ of 10 ROIs in a single field of view was only 0.012 ± 0.005 (mean ± SD) after 30 seconds, suggesting nIROT-SELEC has faster binding dynamics for oxytocin over vasopressin when immobilized. nIROT-SELEC maintained its fluorescence over continuous laser illumination over the course of the 270 second experiment without photobleaching, suggesting that nIROT-SELEC can bind and optically respond to oxytocin without signal attenuation on solid substrates. Importantly, these imaging results suggest that nIROT-SELEC demonstrates full selectivity for oxytocin over vasopressin when used for imaging experiments, enabling its use as an oxytocin nanosensor for brain neurochemical imaging.

Next, to explore nIROT-SELEC compatibility with pharmacology, the nanosensor’s optical response was also tested for a response to oxytocin receptor-targeting drugs. nIROT-SELEC fluorescence modulation was minimal upon treatment with oxytocin receptor agonists atosiban [ΔF/F_0_ = −0.023 ± 0.024 (mean ± SD); n = 3] and L-368,899 [ΔF/F_0_ = −0.16 ± 0.03 (mean ± SD); n = 3], and oxytocin analog and receptor agonist, carbetocin [ΔF/F_0_ = 0.005 ± 0.027 (mean ± SD); n = 3]. These data suggest that our nanosensor can be leveraged to explore potential changes in oxytocin signaling induced by – and in conjunction with – pharmacological agents.

### *Ex vivo* imaging of electrically-evoked oxytocin release in acute brain slices of mice

To explore the use of nIROT-SELEC for *ex vivo* oxytocin imaging, we imaged the nanosensor’s fluorescence response in brain tissue of C57BL/6 mice with the extracellular space labeled with nIROT-SELEC. Acute coronal mouse brain slices of 300 µm thickness were prepared as previously described. ^30,31^ Slices were incubated with 2 mg/L nIROT-SELEC in oxygen-saturated artificial cerebrospinal fluid (aCSF) for 15 min to enable nanosensors to diffuse into the brain tissue, resulting in even and widespread nIROT-SELEC localization in the extracellular space ^21,32^. We subsequently rinsed the slices to remove excess nIROT-SELEC and transferred them to an aCSF-perfused chamber for 15 min before imaging. nIROT-SELEC-labeled slices were imaged using a custom-built nIR upright epifluorescence microscope that excites nanosensors with 785 nm laser and obtains fluorescence from 900 to 1700 nm with an nIR camera with an imaging field of view measuring 175 µm by 140 µm.

We first investigated terminal release of oxytocin in the paraventricular nucleus of hypothalamus (Fig. 4a), where oxytocin is synthesized and somatodendritically released ^1,33^. Applying a 0.5 mA current with a bipolar stimulating electrode evoked an instantaneous increase in nIROT-SELEC fluorescence [ΔF/F_0_ = 0.035 ± 0.007 (mean ± SD); n = 3] as shown in Fig. 4b. Note that the electrical stimulation of neurons does not provide cell specificity, potentially leading to the release of other neurochemicals near the electrodes. However, oxytocin selectivity can be achieved by bath-application of pharmacological agents, which will be discussed later. The fluorescent change in nIROT-SELEC over the entire field of view provides the integrated ΔF/F_0_. To obtain subcellular information, as described in detail elsewhere ^24,31,34^, a 6.8 µm by 6.8 µm grid mask was applied, and the ΔF/F_0_ for each grid square was calculated. A grid square was identified as an ROI if the fluorescence upon stimulation was significant compared to the baseline fluorescence (Fig. 4b and S5) (see Methods for definition of significance). We approximate ROI values to represent the number of oxytocin release sites, and the ΔF/F0 of each release site as a proxy for the amount of oxytocin released per release site. Subsequently, the ΔF/F_0_ contribution from all release sites was averaged over all identified release sites to provide a mean ΔF/F_0_ across release sites (Fig. 4c). This data analysis approach allowed us to disentangle the integrated ΔF/F_0_ into the number of oxytocin release sites and The oxytocin released per release site, enabled by the high spatiotemporal resolution of nIROT-SELEC. To verify their reversibility, we applied three repeated stimulations, 25 seconds apart to enable the tissue to recover. Each stimulation resulted in increased fluorescence [integrated ΔF/F_0_ = 0.023 ± 0.008, 0.020 ± 0.006 and 0.019 ± 0.007 (means ± SD) for the 1st, 2nd, and 3rd stimulation, respectively; n = 2] (Fig. S6), demonstrating nIROT-SELEC’s reversibility.

**Figure 4.**
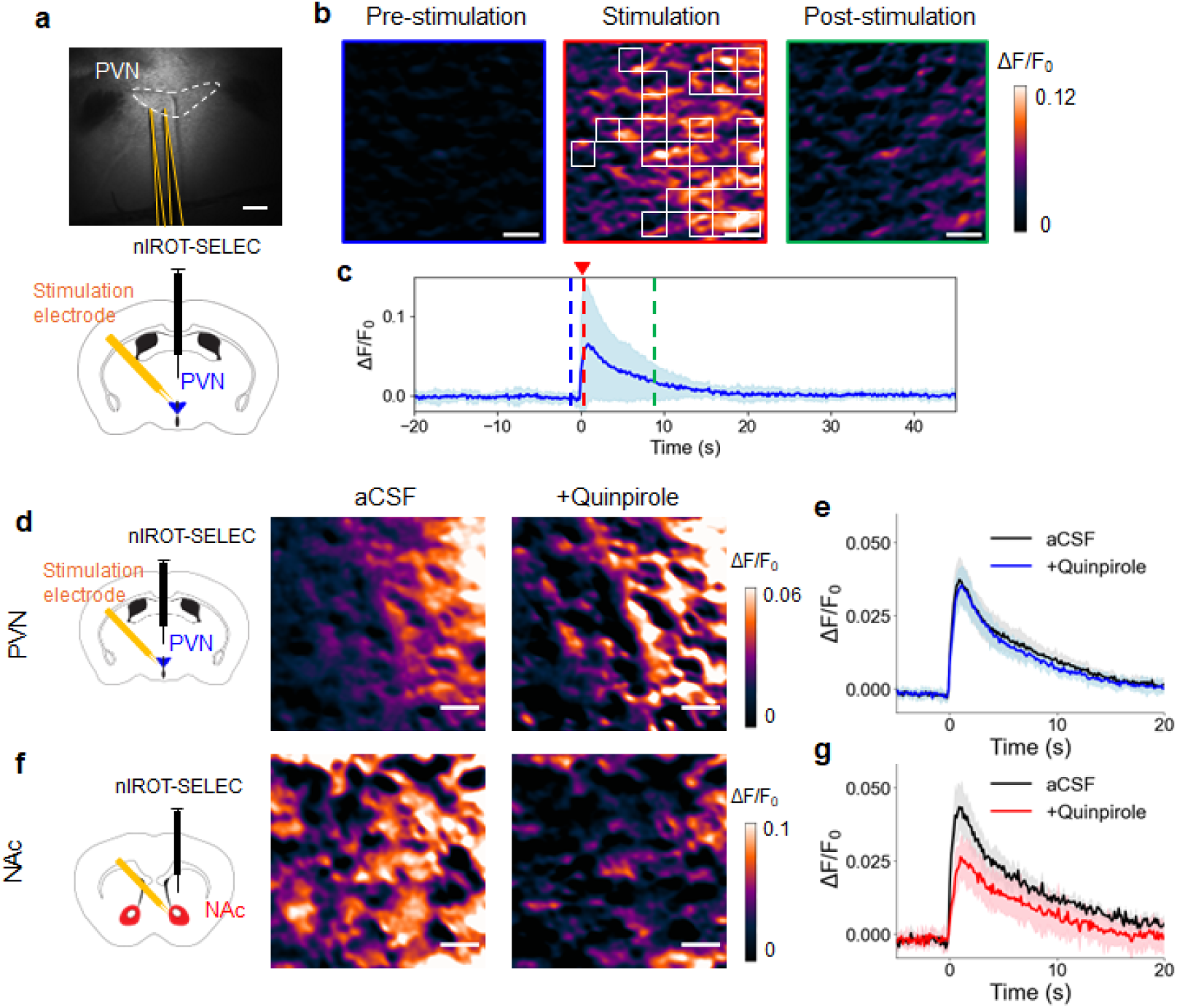
nIROT-SELEC detects neuromodulators *ex vivo* in mice. **(a)** (top) Bright field image and (bottom) schematic image showing the paraventricular nucleus (PVN) in the brain slices with stimulation electrodes (yellow). Scale bar represents 200 µm. **(b)** nIROT-SELEC ΔF/F_0_ images of PVN following 0.5 mA electrical stimulation in standard aCSF. Three frames are shown: (left) “pre-stimulation” is the baseline ΔF/F_0_ before electrical stimulation, (center) “stimulation” is immediately after electrical stimulation, where a 6.8 µm by 6.8 µm grid mask was applied and grids with high ΔF/F_0_ were identified as ROIs (white), and (right) “post-stimulation” is after ΔF/F_0_ has mostly returned to baseline. Scale bars represent 10 µm. Note that grid masks here are only for schematic purposes and do not represent actual grids identified by the analysis. **(c)** Time-course ΔF/F_0_ averaged over ROIs in a single field of view (175 µm by 140 µm) in the PVN. The solid line is the averaged value over identified ROIs, with the shaded region representing standard deviation. Blue, red, and green dashed lines represent where the pre-stimulation, stimulation, and post-stimulation images in (b) are taken from, respectively. **(d)** ΔF/F_0_ images of PVN after stimulation (left) before and (right) after 2 µM quinpirole addition. Scale bars represent 10 µm. **(e)** Time-course integrated ΔF/F_0_ with (red) and without (black) 2 µM quinpirole addition in the PVN. The solid line is the averaged value over three animals, with the shaded region representing standard deviation. **(f)** ΔF/F_0_ images of nucleus accumbens (NAc) after stimulation (left) before and (right) after 2 µM quinpirole addition. Scale bars represent 10 µm. **(g)** Time-course integrated ΔF/F_0_ with (red) and without (black) 2 µM quinpirole addition in the PVN. The solid line is the averaged value over three animals, with the shaded region representing standard deviation.

Since nIROT-SELEC responds to dopamine (Fig. 3c), to confirm the selectivity of oxytocin release in brain slice imaging experiments, we sought to bath-apply quinpirole, a D2 receptor agonist known to suppress dopamine release. First, we conducted *in vitro* solution phase experiments (without biological tissue) to test the nIROT-SELEC fluorescent response towards quinpirole. Application of 2 µM quinpirole did not cause any fluorescent change to nIROT-SELEC [ΔF/F_0_ = −0.06 ± 0.03 (means ± SD); n = 3] (Fig. S7), confirming that quinpirole can be used concurrently with nIROT-SELEC to suppress dopamine release for selective oxytocin imaging in brain tissue. Furthermore, the nanosensor *in vitro* ΔF/F_0_ response to oxytocin was not altered by simultaneous application of 2 µM quinpirole [ΔF/F_0_ = 1.00 ± 0.10 for oxytocin only and ΔF/F_0_ = 0.91 ± 0.04 for oxytocin and quinpirole (means ± SD); *P* = 0.237; n = 3] (Fig. S7), confirming that nIROT-SELEC retained their functionality in the presence of quinpirole. We next used 2 µM quinpirole to suppress dopamine release in brain slices containing two regions: the paraventricular nucleus of hypothalamus, where a limited amount of dopamine was previously detected ^35,36^, and the nucleus accumbens, in which oxytocin signaling is of great interest to elucidate social behaviors including bond formation ^37^ and social reward ^38^, but where higher dopamine concentration is expected ^39–41^. Although quinpirole application did not cause a change in nIROT-SELEC fluorescence intensity when imaging in the paraventricular nucleus of hypothalamus relative to quinpirole-free slice imaging [integrated ΔF/F_0_ = 0.035 ± 0.007 in aCSF and integrated ΔF/F_0_ = 0.034 ± 0.005 in aCSF and quinpirole (means ± SD); n = 3; *P* = 0.951] (Fig. 4d and e), quinpirole reduced the nanosensor fluorescence intensity by 42% in the nucleus accumbens [integrated ΔF/F_0_ = 0.042 ± 0.007 in aCSF and integrated ΔF/F_0_ = 0.024 ± 0.006 in aCSF and quinpirole (means ± SD); n = 3; *P* = 0.016] (Fig. 4f and g), confirming the efficacy of quinpirole application. This observation also suggests that we measure lower dopamine concentrations in the paraventricular nucleus of hypothalamus than those in the nucleus accumbens, agreeing with previous microdialysis studies in the paraventricular nucleus of hypothalamus ^35,36^ and in the nucleus accumbens ^39–41^. While the integrated ΔF/F_0_ did not decrease by quinpirole application in the paraventricular nucleus of hypothalamus, the number of oxytocin release sites decreased by 22% in the paraventricular nucleus of hypothalamus [release sites = 290 ± 30 in aCSF and release sites = 225 ± 66 in aCSF and quinpirole (means ± SD); n = 3; *P* = 0.127] and by 40% in the nucleus accumbens [release sites = 321 ± 59 in aCSF and release sites = 193 ± 65 in aCSF and quinpirole (means ± SD); n = 3; *P* = 0.0988] (Fig. S8). Since release-site analysis is more sensitive than overall changes in nanosensor fluorescence ^34^, we predict that a small number of dopamine release sites are present in the paraventricular nucleus of hypothalamus, and a larger number of dopamine release sites are present in the nucleus accumbens, which are suppressed by quinpirole.

### Application of nIROT-SELEC to prairie voles for *ex vivo* imaging of oxytocin signaling

One advantage of using non-genetically encoded sensors like nIROT-SELEC is their applicability to comparative neuroscience, including their use to study non-traditional model organisms. To demonstrate this versatility, we imaged evoked release of neurochemicals in prairie voles (*Microtus ochrogaster*) - a rodent species that exhibits the unusual traits of social monogamy and selective affiliation for familiar conspecifics. ^42^ While rats and mice are commonly studied in neuroscience, they are socially gregarious and do not display selective preferences for familiar peers or mates. ^43–45^ Like prairie voles, humans form selective relationships, ^46,47^ thus understanding the oxytocin function in prairie voles’ selective affiliations is of great interest and importance to extrapolate the role of oxytocin in human peer interactions.

We first imaged brain slices of prairie voles containing the nucleus accumbens (Fig. 5a), a region associated with bond formation ^37^, social reward ^38^, maternal interaction ^48^, and selective affiliation ^6,13^. Brain slices were prepared and incubated with nIROT-SELEC as described above for mice, which demonstrates the ease of preparation and translation to another species. Upon electrical stimulation, we observed an increase in nIROT-SELEC fluorescence [integrated ΔF/F_0_ = 0.033 ± 0.015 (mean ± SD); n = 9] (Fig. 5a and b), confirming nIROT-SELEC’s applicability to prairie voles. As electrical stimulation results in off-target neurochemical release, quinpirole was bath-applied to suppress dopamine release, which decreased nIROT-SELEC fluorescence intensity by 33% [integrated ΔF/F_0_ = 0.022 ± 0.011 in aCSF and quinpirole (means ± SD); n = 9; *P* = 0.0354] (Fig. 5a and b). Release site analysis revealed a 72% reduction in the number of release sites following quinpirole application [ROIs = 372 ± 59 in aCSF and release sites = 268 ± 75 in aCSF and quinpirole (means ± SD); n = 9; *P* = 0.0082], while the mean ΔF/F_0_ across release sites remained at 98% (Fig. 5c) [mean ΔF/F_0_ = 0.042 ± 0.017 in aCSF and integrated ΔF/F_0_ = 0.041 ± 0.021 in aCSF and quinpirole (means ± SD); n = 9; *P* = 0.8203], consistent with observations in mice and in other studies ^24,34^ that suggest quinpirole affects the number of release sites more than the mean ΔF/F_0_ across release sites.

**Figure 5.**
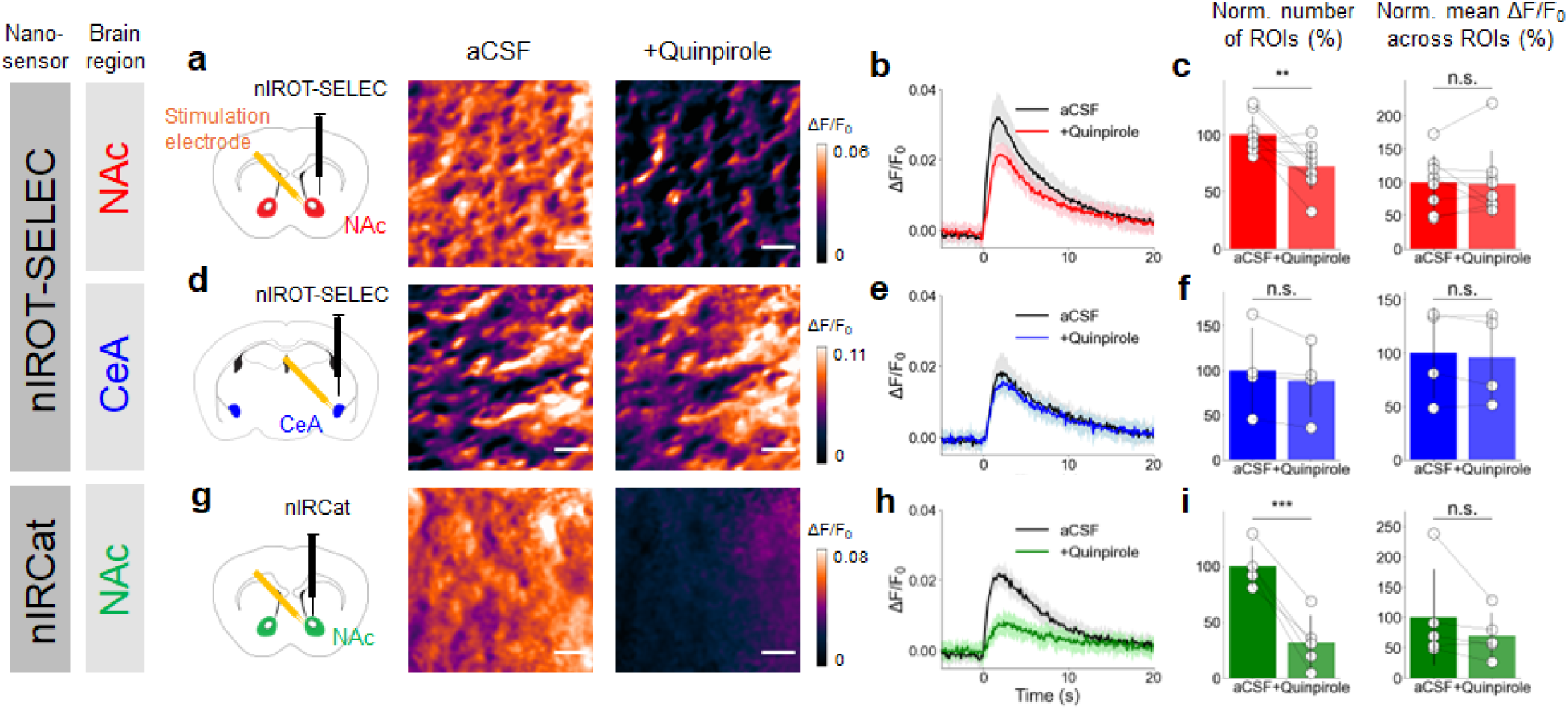
nIROT-SELEC detects neuromodulators *ex vivo* in prairie voles. **(a-c)** nIROT-SELEC applied to the nucleus accumbens (NAc). **(a)** ΔF/F_0_ images of nucleus accumbens after stimulation (left) before and (right) after 2 µM quinpirole addition. Scale bars represent 10 µm. **(b)** Time-course integrated ΔF/F_0_ with (red) and without (black) 2 µM quinpirole addition in the nucleus accumbens. The solid line is the averaged value over nine animals, with shadow representing SD. **(c)** (left) Normalized number of ROIs in aCSF and in aCSF with quinpirole. Each datapoint is normalized to the average value of ROIs in aCSF. Error bars represent SD [n = 9; *P* = 0.0082]. (right) Normalized mean ΔF/F_0_ across ROIs in aCSF and in aCSF with quinpirole. Each datapoint is normalized to the average value of mean ΔF/F_0_ in aCSF. Error bars represent SD [n = 9; *P* = 0.8203]. **(d-f)** nIROT-SELEC applied to the central nucleus of the amygdala (CeA). **(d)** ΔF/F_0_ images of CeA after stimulation (left) before and (right) after 2 µM quinpirole addition. Scale bars represent 10 µm. **(e)** Time-course integrated ΔF/F_0_ with (blue) and without (black) 2 µM quinpirole addition in the CeA. The solid line is the averaged value over four animals, with shadow representing SD. **(f)** (left) Normalized number of release sites in aCSF and in aCSF with quinpirole. Each datapoint is normalized to the average value of release sites in aCSF. Error bars represent SD [n = 4; *P* = 0.1635]. (right) Normalized mean ΔF/F_0_ across release sites in aCSF and in aCSF with quinpirole. Each datapoint is normalized to the average value of mean ΔF/F_0_ in aCSF. Error bars represent SD [n = 4; *P* = 0.3515]. **(g-i)** nIRCat applied to the nucleus accumbens. **(g)** ΔF/F_0_ images of nucleus accumbens after stimulation (left) before and (right) after 2 µM quinpirole addition. Scale bars represent 10 µm. **(h)** Time-course integrated ΔF/F_0_ with (green) and without (black) 2 µM quinpirole addition in the nucleus accumbens. The solid line is the averaged value over five animals, with shadow representing SD. **(i)** (left) Normalized number of release sites in aCSF and in aCSF with quinpirole. Each datapoint is normalized to the average value of release sites in aCSF. Error bars represent SD [n = 5; *P* = 0.0008]. (right) Normalized mean ΔF/F_0_ across release sites in aCSF and in aCSF with quinpirole. Each datapoint is normalized to the average value of mean ΔF/F_0_ in aCSF. Error bars represent SD [n = 5; *P* = 0.1250]. Absolute values for (c), (f), (i) are available in Fig. S9. Statistical significance annotation follows *** for *p* < 0.001, ** for *p* < 0.01, * for *p* < 0.05, and n.s. for p > 0.05.

Next, we applied nIROT-SELEC to brain slices containing the central nucleus of the amygdala (Fig. 5d), a region implied in social recognition, memory, and salience ^13,48^. In the central nucleus of the amygdala, the observed integrated ΔF/F_0_ was smaller than that in the nucleus accumbens [integrated ΔF/F_0_ = 0.018 ± 0.011 (mean ± SD); n = 4], with only an 11% decrease following quinpirole application (Fig. 5e and f) [integrated ΔF/F_0_ = 0.016 ± 0.010 in aCSF and quinpirole (means ± SD); n = 4; *P* = 0.0335], consistent with previous studies detecting limited dopamine concentration in the central nucleus of the amygdala ^49,50^. Neither the number of release sites, nor mean ΔF/F_0_ across release sites showed statistically significant changes with quinpirole application within the central nucleus of the amygdala, as expected (Fig. S9).

It is possible that the remaining ΔF/F_0_ in the nucleus accumbens in Fig. 5a-c may be from dopamine not fully suppressed by quinpirole. To test this, we applied nIRCat, a near infrared catecholamine nanosensor for detecting dopamine ^21^, instead of nIROT-SELEC to the nucleus accumbens. Electrical stimulations increased nIRCat fluorescence [integrated ΔF/F_0_ = 0.022 ± 0.018 in aCSF (mean ± SD); n = 5] (Fig. 5g and h), consistent with a previous study ^21^. Notably, integrated ΔF/F_0_ after quinpirole application was 36% of that before application [integrated ΔF/F_0_ = 0.008 ± 0.006 in aCSF and quinpirole (means ± SD); n = 5; *P* = 0.0625] (Fig. 5i), contrasting with the 67% observed with nIROT-SELEC in the nucleus accumbens (Fig. 5c). This suggests that if all signals observed in the nucleus accumbens with nIROT-SELEC (Fig. 5a-c) were attributed to dopamine, the expected suppression by quinpirole would be around 64%, but the observed reduction was 33%. Thus, the dataset with nIRCat in the nucleus accumbens strongly suggests that nIROT-SELEC with quinpirole is capable of detecting non-dopamine signaling, i.e., oxytocin.

## Discussion

Sensitive and selective high spatiotemporal imaging of neuropeptides is crucial to understanding the neurochemistry underpinning social behavior. Synthetic optical probes based on carbon nanotubes offer a promising approach to imaging neuropeptide signaling with numerous advantages, including their compatibility with pharmacology and species- and age-independent application. Here, we demonstrate a method for evolving synthetic neuropeptide nanosensors with SELEC, and demonstrate the use of this evolution method to develop an oxytocin nanosensor for oxytocin imaging in living brain tissue. We identified a nanosensor, termed nIROT-SELEC, with a ΔF/F_0_ of 1.72±0.01 upon addition of 100 µM oxytocin. We showed that nIROT-SELEC has a K_d_ for oxytocin of 4.93±3.19 µM. Further, we demonstrated that nIROT-SELEC is selective for oxytocin over oxytocin analogs, receptor agonists and antagonists, and most other neurochemicals. Together, these data demonstrate that SELEC can be used to achieve rapid identification of ssDNA-SWCNT nanosensors for neuropeptide targets, which are larger and more structurally complex than most neurochemicals, potentiating the use of SELEC as a generic method to evolve synthetic nanosensors for diverse neuropeptide imaging applications across species.

Excitingly, for the first time, the non-genetically encoded nature of nIROT-SELEC enabled real-time imaging neurochemical signaling in prairie voles, a rodent species used prolifically in social neuroscience because of their propensity to form selective social bonds. A comparative approach and the use of appropriate animal models is crucial, especially in social neuroscience, because the forms of social behaviors displayed by species are diverse. For instance, while humans are typically considered to display social monogamy ^46,47^, this trait is very rare in mammals, observed in only 9% of species ^51^. Thus the neural basis of social monogamy and its health consequences requires the use of non-traditional animals like prairie voles, as model organisms like mice or rats typically prefer social novelty and do not form social bonds ^44,45^. Furthermore, studying diverse species is essential to uncover the variety of pathways supporting behaviors, as well as the generalizability or translatability of findings across species ^52–54^. Comparative studies will eventually enable us to understand the evolutionary origins of these neural pathways and offer flexibility in selecting research organisms that display human-like behaviors. While it is possible to develop genetically encoded sensors for each species, purely synthetic sensors like nIROT-SELEC offer a more efficient solution for cross-species application once established, while the synaptic-scale imaging resolution of SELEC-based probes offers a more robust metric for comparison across species and phenotypes than arbitrary unit fluorescence modulation. Such techniques open the door to a variety of models where neurochemistry has not yet been thoroughly studied, potentially holding the key to understanding complex neural pathways and neurological disorders.

Although nIROT-SELEC, similarly to most ssDNA-SWCNT, displayed non-specific fluorescence modulation (normalized ΔF/F_0_ = 1.55 ± 0.26) to dopamine, we demonstrate that quinpirole can be used to suppress dopamine release for selective oxytocin imaging in brain tissue. Application of quinpirole effectively suppressed dopamine release in *ex vivo* measurements (Fig. 4 and 5), as the synthetic nature of nIROT-SELEC enables concurrent use of pharmacological agents during brain slice imaging. Furthermore, the unique cross-sensitivity of nIROT-SELEC for both oxytocin and dopamine can be used to provide insights on the role and relative presence of both neuromodulators and the relative density of their release sites in tissue. In this work, we demonstrated the ability to do so by imaging oxytocin and dopamine release simultaneously, and next comparing these data to tissues exposed to D2 receptor agonists to calculate the relative contributions of dopamine, and dopamine release sites, in brain regions where both neuromodulators play a central neurobiological role. For example, the number of release sites was 321 ± 59 (means ± SD) in nucleus accumbens before quinpirole application and was reduced to 193 ± 65 (means ± SD) after quinpirole application (n = 3). The quinpirole-specific reduction in neuromodulator release sites (release sites = 128) can be attributed to endogenous dopamine release sites, and the release sites that persist in the presence of quinpirole (release sites = 193) correspond to oxytocin release, providing information about *both* neurochemicals with a *single* nanosensor. nIROT-SELEC could thus serve as an efficient platform to study the interplay between oxytocin and dopamine, which has become of great interest due to the hypothesized interplay of dopamine and oxytocin for maternal interaction, social reward, and pair-bond formation ^48,55–57^. nIROT-SELEC also responds moderately to vasopressin with a normalized ΔF/F_0_ = 0.69 ± 0.25 over a 20-minute long solution-phase incubation *in vitro*, but immobilizing nIROT-SELEC for imaging-based fluorescence quantification provides selectivity for oxytocin over vasopressin. nIROT-SELEC also responds minimally to oxytocin receptor agonists and antagonists, indicating its potential to study the effects of receptor agonists and antagonists such as atosiban, carbetocin, and L-368,899 in the extracellular space.

To the best of our knowledge, real-time imaging of oxytocin in non-traditional animals has not been demonstrated, making our *ex vivo* brain slice imaging of oxytocin a promising achievement. This technique can provide unprecedented molecular and neurochemical insights into animal behaviors (e.g., social monogamy) in the future. Nonetheless, achieving *in vivo* imaging is a crucial future step, as it would enable us to monitor oxytocin activity in the awake and behaving brain. Such techniques would allow us to understand timing and location of the oxytocin release in relation to complex animal behaviors. *In vivo* imaging has been largely accomplished with fiber photometry-based approaches in conjunction with genetically encoded probes, but these approaches eliminate access to sub-cellular spatial information that is critical for understanding neurochemical signaling. Efforts towards *in vivo* imaging, including fiber photometry ^58^ and dual near infrared microscopy ^59,60^ are underway, with the former compromising on the unique spatial information achievable with SELEC-based probes. As such, deep-tissue imaging is particularly promising, as the unique near infrared fluorescence of nIROT-SELEC might circumvent the need for surgical windows and enable the concurrent study of animal behavior with sub-cellular resolution neurochemical imaging. Another important application of oxytocin imaging will be its use across developmental stages. The oxytocin system undergoes drastic changes during development and is believed to be both a target and mediator of early life experience ^61^. Genetically encoded sensors typically require weeks to express, leaving the dynamics and effects of oxytocin during development largely unknown. Since nIROT-SELEC does not require genetic manipulation, it is readily transferable to imaging oxytocin signaling in developing brains.

In summary, we developed a method to evolve molecular recognition for neuropeptides, and demonstrate the feasibility of our method to develop a neuropeptide for oxytocin, nIROT-SELEC. Our oxytocin probe provides an unprecedented ability to study oxytocin across species and developmental stages, with spatial resolution that remains unachievable with other techniques. Ultimately, our results suggest that nIROT-SELEC nanosensors offer a novel method for high spatiotemporal fluorescence imaging to study oxytocin signaling in the brain, to elucidate the role of oxytocin in both health and disease.

## Materials and Methods

### Reagents

Small diameter HiPCo single-walled carbon nanotubes (SWCNT) were purchased from NanoIntegris. Oxytocin acetate salt hydrate, [Arg8]-vasopressin acetate salt, dopamine hydrochloride, thyrotropin-releasing hormone (TRH), glutamate, γ-aminobutyric acid, and atosiban were purchased from Millipore Sigma. Quinpirole and carbetocin were purchased from Tocris Bioscience. ssDNA sequences were purchased from Integrated DNA Technologies.

### SELEC experimental protocol

The SELEC protocol was adapted from a previous paper.^19^ The control sequence library from Jeong et al. was preserved. A new control sequence library (evolved in the absence of a target analyte) was thus not developed in this study.

The initial ssDNA library used for evolution consisted of 18 random nucleotides flanked by 1) two 6-mer polycytosines and 2) two 18-mer primer regions for polymerase chain reaction: AGCGTCGAATACCACTAC-C_6_-N_18_-C_6_-GACCACGAGCTCCATTAG. During the first round of evolution, 100 nmol of the ssDNA library (200 μL, 0.5 mM) was mixed with pH 5 acetate buffer (200 μL, 20 mM) and 10 μg of SWCNT. The mixture was bath sonicated for 5 min, heated at 95°C for 5 min to denature the ssDNA, then cooled to room temperature. Next, 100 µL of 1 mM oxytocin was added for a final volume of 500 μL. The mixture was allowed to incubate for 10 min at room temperature prior to probe tip sonication (Cole-Parmer Ultrasonic Processor, CPX130) with a 3 mm tip in ice-cold water for 20 min at 30% amplitude. The resulting suspension was centrifuged for 60 min at 16,000 g to pellet unsuspended SWCNT. The supernatant containing ssDNA-SWCNT and free ssDNA was collected and spin-filtered in a 0.5 mL 100 kDa MWCO Amicon Ultra cassette at 5,000 g for 2 min six times with DNAse-free water to remove unbound ssDNA. The suspension was then mixed with 0.1% (w/v) SDBS in 1X PBS (200 μL) and heated at 95°C for 1 hr to detach bound ssDNA from the SWCNT surface. The suspension was then centrifuged for 60 min at 16,000 g to pellet aggregated SWCNT, and the supernatant containing ssDNA was collected. The solution was spin-filtered in a 0.5 mL 3 kDa MWCO Amicon Ultra cassette four times with DNAse-free water to remove SDBS.

The collected ssDNA was amplified by PCR, using a previously described protocol, with a FAM-modified forward primer (FAM-AGCGTCGAATACCACTAC) and biotinylated backward primer (biotin-CTAATGGAGCTCGTGGTC). The following PCR mixture was prepared for both preliminary and preparative PCR: Hot Start Taq DNA polymerase (2.75 U, New England Biolabs), 1X Hot Start Taq reaction buffer, forward primer (1 μM), backward primer (1 μM), deoxynucleotide triphosphate (500 μM), and ssDNA library template (∼100 ng/mL). The total PCR mixture volume for each preliminary PCR reaction tube and preparative PCR well was 100 μL. DNAse-free water was added to achieve the desired volume. The following standard cycling conditions were used: initial denaturation for 900 s at 95°C, *N* cycles of denaturation for 30 s at 95°C, annealing for 30 s at 50°C, and extension for 30 s at 72°C, and final extension for 180 s at 72°C.

Preliminary PCR was used to determine the number of PCR cycles (*N*) necessary to maximize dsDNA yield and minimize PCR by-product formation. Two PCR reaction tubes, with and without the ssDNA library template, were collected at the following cycles: 10, 15, 20, 25 and 30. A 4% agarose gel was prepared by heating 30 mL of 1X TBE buffer and 1.2 g of low range ultra agarose (Bio-Rad Laboratories) in a microwave oven and electrophoresis was performed on the preliminary PCR samples at 110 V for 18 min. The gel was stained with SYBR Gold, and the DNA bands were observed under UV light. The cycle number that yielded the brightest DNA band without nonspecific amplicons was selected for preparative PCR.

For preparative PCR, a 10 mL PCR mixture was prepared to produce a large quantity of ssDNA for the subsequent round of evolution. 100 μL of the PCR mixture was added to 95 of 96 reaction wells. A negative control well was prepared without the ssDNA library template. The preparative PCR product from a single reaction well was collected, purified with a GeneJet PCR purification kit (Thermo Scientific), and stored at −20°C for sequencing preparation. The remaining preparative PCR product was spin-filtered using the 15 mL 10 kDa MWCO Amicon Ultra centrifugal filter.

To extract amplified ssDNA from the PCR product, 2.5 mL of Streptavidin-Sepharose beads (GE Healthcare) were first added to a mounted sintered Buchner funnel (<10 μm pore size). The beads were washed with 10 mL PBS buffer. PCR product was then incubated with beads for 30 min and passed through the funnel 3 times upon which the beads were washed again with 10 mL PBS. To elute and collect FAM-labeled ssDNA, 8 mL 0.2 M NaOH aqueous solution was added slowly to the beads. To desalt the ssDNA, an NAP-10 desalting column (GE Healthcare) was washed with deionized water (20 mL). 1 mL ssDNA solution was then added and drained. Upon the addition of 1.5 mL DNAse-free water, the eluate was collected. Desalted ssDNA was concentrated by spin-filtering in a 15 mL 3 kDa MWCO Amicon Ultra centrifugal filter and dried using a DNA Speedvac. For subsequent rounds of SELEC, the dried ssDNA was reconstituted in pH 5 acetate buffer, and the concentration was determined by measuring the absorbance at 260 nm.

### High-throughput sequencing and analysis

To achieved multiplexed sequencing of SELEC libraries, two sequential PCR steps were used to add Illumina TruSeq universal adapter sequences and sequencing indices. Libraries from SELEC rounds 3 to 6 were sequenced with an Illumina HiSeq 4000 at the Genomic Sequencing Laboratory at the University of California, Berkeley. From the ∼20 million raw sequences generated from each round, we discarded sequences that lacked the correct fixed regions (the 18-mer PCR primer regions and C_6_ anchor regions). The FASTAptamer toolkit was used to filter and count sequence frequencies. All experiments in this manuscript used ssDNA sequences which excluded the 18-mer PCR primer regions. These sequences thus only contained the 18-mer random region flanked by the 6-mer polycytosine regions (i.e. C_6_-N_18_-C_6_). Data visualizations of the top 200 sequences from the experimental and control SELEC libraries were performed using matplotlib and MATLAB.

### Synthesis of ssDNA-SWCNT nanosensors

ssDNA-SWCNT nanosensors were prepared in a 1.5 mL DNA LoBind tube to a final volume of 1 mL by combining HiPCo SWCNT slurry (900 µL, 0.22 mg/mL in 1X PBS) and ssDNA (100 µL, 1 mM in H_2_O). SWCNT slurry was vortexed for 3 sec and bath sonicated for 10 min prior to use. ssDNA were heated at 55°C for 5 min and cooled to room temperature before use. The mixture was vortexed for 3 sec, bath sonicated for 10 min, and probe-tip sonicated (Cole-Parmer Ultrasonic Processor, 3-mm tip) in ice-cold water for 10 min at 50% amplitude. The resulting suspensions were centrifuged at 16,000 g for 60 min to pellet unsuspended SWCNT. 900 µL of the supernatant was centrifuged again for 30 min. From this, 850 µL of the supernatant was carefully extracted and transferred to a new tube. Nanosensor concentration was calculated by measuring absorbance at 632 nm (NanoDrop One, Thermo Scientific) with an extinction coefficient of ε = 0.036 (mg/L)^-1^ cm^-1^. Nanosensors were diluted to 5 mg/L in PBS for characterization experiments and were allowed to equilibrate at 4°C for at least 24 hours prior to use in experiments.

### Optical characterization of ssDNA-SWCNT nanosensors

Near-infrared fluorescence spectra were collected using a custom-built spectrometer and microscope as described previously^62^. Measurements were obtained with a 20X objective on an inverted Zeiss microscope (Axio Observer.D1) coupled to a spectrometer (SCT 320, Princeton Instruments) and liquid nitrogen cooled InGaAs linear array detector (PyLoN-IR, Princeton Instruments). Nanosensor suspensions were excited with a 721 nm laser (OptoEngine LLC) inside a polypropylene 384 well plate (Greiner Bio-One microplate). Spectra were collected with a 500 ms integration time.

For analyte screening, the baseline near-infrared fluorescence spectrum of each nanosensor-containing well was collected. PBS or analyte in PBS were added to wells, and fluorescence spectra were collected at regular time points until the maximum analyte fluorescence response was achieved (∼20 min following addition). Responses were calculated and reported as ΔF/F_0_ = (F_t_-F_0_)/ F_0_, where F_0_ is the baseline nanosensor fluorescence and F_t_ is the peak fluorescence after analyte incubation at timepoint *t*. The peak fluorescence corresponds to the (8,6) SWCNT chirality, which has a maximum near-infrared fluorescence at ∼1195 nm. To correct for variable fluctuations in the fluorescence measurement over time, the ΔF/F_0_ of the PBS control was subtracted from the analyte ΔF/F_0_.

Surface-immobilized nanosensors were imaged on an epifluorescence microscope (100X oil immersion objective) and a Ninox VIS-SWIR 640 camera (Raptor) and excited with a 721 nm laser. For each imaging experiment, 120 μL PBS was added and the z-plane was refocused prior to recording. Image stacks were collected with a 950 ms exposure time and 1 Hz frame rate for 270 seconds and total frames.

For immobilization experiments, nanosensors were immobilized on MatTek glass-bottom microwell dishes (35 mm petri dish with 10 mm microwell). The dish was washed twice with 1X PBS (150 μL), 15 mg/L nanosensors (100 μL, in PBS) were added, incubated for 30 min, and removed, then the dish was washed twice again with 1X PBS (150 μL). For imaging immobilized SWCNT, PBS was added at frame 60 and analyte was added at frame 120.

Image stacks were processed in ImageJ by applying a median filter (2.0-pixel radius) and gaussian filter (2.0-pixel radius). ROI analysis was automated using a Python script. ROI size was set to 25×25 pixels. Responses were calculated as ΔF/F_0_ = (F_f_-F_0_)/ F_0_ for each ROI, where F_0_ was the mean integrated fluorescence across the ROI at frame 1 and F_f_ was the mean integrated fluorescence for frame *f*, where *f* = 1-300. To account for microscope drift, slope was calculated from frame 1 to frame 110 using the sklearn.linear_model Linear Regression function and used to determine and subtract drift from the overall integrated fluorescence. ΔF/F_0_ for the top 10 ROIs (ranked by response at *f* = 300) were averaged and presented with standard deviation.

### Acute slice preparation and nanosensor labeling

C57BL/6 strain male mice were purchased from Jackson Laboratory (stock no. 000664) and were 14-17 weeks old when imaged. The mice were weaned on postnatal day 21, and group-housed with nesting material on a 12:12 light cycle. All animal procedures were approved by the UC Berkeley Animal Care and Use Committee (ACUC). Prairie voles were bred from in-house colonies in a long photoperiod (14 hr light:10 hr dark; 14 h light; 04:00 to 18:00 PST). Voles were weaned at postnatal day 21 and separated into same-sex pair-housing in clear plastic cages with sani-chip bedding and an opaque plastic hiding tube. All procedures adhered to federal and institutional guidelines and were approved by the Institutional Animal Care and Use Committee at the University of California, Berkeley.

Acute brain slices were prepared following the protocols described in Ref. ^30,31^. Cutting buffer (119 mM NaCl, 26.2 mM NaHCO_3_, 2.5 mM KCl, 1mM NaH_2_PO_4_, 3.5 mM MgCl4, 10 mM glucose, all purchased from Sigma-Aldrich) and aCSF (119 mM NaCl, 26.2 mM NaHCO_3_, 2.5 mM KCl, 1mM NaH_2_PO_4_, 1.3 mM MgCl_2_, 10 mM glucose, 2 mM CaCl_2_, all purchased from Sigma-Aldrich) were prepared and bubbled with carbogen gas (oxygen/carbon dioxide 95% O_2_, 5% CO_2_, Praxair, cat. no. BIOXCD5C-K). The animals were deeply anesthetized via intraperitoneal injection of ketamine (120 mg/kg) and xylazine (24 mg/kg) and transcardially perfused with 5 mL of ice-cold cutting buffer. The brain was subsequently extracted, mounted on a vibratome cutting stage (Leica VT 1000) in the carbogen-bubbled cutting buffer, and cut into 300 µm thick coronal slices with microtome razor blades (Personna, double edge super blades; VWR, cat. no. 100491-888). Slices were incubated at 37°C for 30 min in oxygen saturated aCSF in a chamber (Scientific Systems Design, Inc., BSK4) and then transferred to room temperature for 30 min. To label the slice with nanosensors, slices were transferred to a separate incubation chamber (Scientific Systems Design, Inc., BSK2) filled with 5 mL of carbogen-bubbled aCSF at room temperature. Nanosensors were applied to the surface of brain slices with a pipette to a final concentration of 2 mg/L and incubated for 15 min. Slices were rinsed for 5 sec with bubbled aCSF through 3 wells of a 24-well plate to remove unlocalized nanosensors, transferred back to the main chamber, and rested for at least 15 min before imaging.

### nIR Microscope Design

A home-built upright epifluorescent microscope (Olympus, Sutter Instruments) was used to image fluorescent response on nanosensors embedded in acute brain slices. 785-nm CW diode laser (OptoEngine LLC, MDL-III-785R-300mW) was used for excitation. After passing through a custom fluorescence filter cube [excitation: 800 nm shortpass (Thorlabs, FESH0800), dichroic: 900 longpass (Thorlabs, DMLP990R), and emission: 900 longpass (Thorlabs, FELH0900)], the beam reaches sample via a 60X objective (Nikon, CFI Apochromat NIR 60X W). Fluorescnece was collected with a two-dimensional InGaAs array detector (Raptor Photonics, Ninox 640). Microscope operation was controlled by Micro-Manager Open Source Microscopy Software ^63^.

### Electrical stimulation evoked neurochemical imaging with nIR microscope

Carbogen-bubbled aCSF was flown through the microscope chamber with a perfusion pump (World Precision Instruments, Peri-Star Pro) and an aspirator pump (Warner Instruments, Dedicated Workstation Vacuum System) at 2 mL/min. The imaging chamber’s temperature was kept at 32°C with an inline heater with feedback control (Warner Instruments, TV-324C). Slices labeled with nanosensors were placed in the chamber with a Tissue harp (Warner Instruments), and a bipolar stimulation electrode (MicroProbes for Life Science Stereotrodes Platinum/Iridium Standard Tip) was positioned to the targeted field of view, which was adjusted with a 4X objective (Olympus XLFluor 4X/340) followed by a 60X objective and was at least 10 µm away from the stimulation electrode. Using Micro-Manager, nIR fluorescent images were acquired at frame rates of 8 frames/s (nominal) for 600 frames, where 1 millisecond of 0.5 mA stimulation was applied at after 200 frames of baseline. This image acquisition was repeated three times with the same field of view, with 5 min of waiting time in between. Subsequently 2 µM of quinpirole (Fisher Scientific, 10-611-0) was applied to the chamber through aCSF perfusion and slices were incubated for 15 min before imaging. Images were acquired in the same field of view as the one before quinpirole applications for three times with 5 min between each stimulation.

### Image processing and data analysis of nanosensor fluorescence response

Imaging movie files were processed using a custom MATLAB application (https://github.com/jtdbod/Nanosensor-Brain-Imaging) or Python code (https://github.com/NicholasOuassil/NanoImgPro) (schematically summarized in Fig. S5). Integrated ΔF/F_0_ (ΔF/F_0_ for the entire field of view, 175 µm by 140 µm) (Fig. 4e, 4g, and S5b) was calculated as ΔF/F_0_ = (F-F_0_) /F_0_, where F_0_ is the average intensity for the first 5% of frames and F is the dynamic fluorescence intensity from the entire field of view and averaged over three stimulation replicates. Next, a 25×25 pixel (corresponding to 6.8 µm by 6.8 µm) grid mask was applied to the image stack to minimize bias and improve stack processing time. For each grid square, a moving average was used to calculate the baseline and subtracted from the original trace. The resulting trace left for the square was a time series of change in fluorescence. Next, between frames 200 and 300 the algorithm looked for an event that was 3 standard deviations above the average noise previously calculated for this square. If an event was found, this square was marked ROI (Fig. S5e). If no event was detected, this square was deemed inactive and no more calculations were completed. Lastly, among identified ROIs, ΔF/F_0_ was averaged over all ROIs, corresponding to the mean ΔF/F_0_ across ROIs (Fig. S5f). The number of ROIs and the mean ΔF/F_0_ of ROIs are averaged over at least three experimental replicates. ΔF/F_0_ for the entire field of view can also be obtained by applying a 640×640 pixel to the Python code above.

## Author contributions

N.N. performed the iterative process of SELEC. N.N. and J.A.M.A. analyzed the high-throughput sequencing data. N.N., J.A.M.A., X.S., E.L., and O.I.A.S characterized the nanosensors *in vitro*. A.K.B. conceptualized the brain regions and A.M.B. raised prairie voles and prepared brain tissue. N.K., A.M.B., and J.Z. performed the *ex vivo* experiments. J.A.M.A., N.K., N.N. and M.P.L. wrote the manuscript. All authors discussed the results and commented on the manuscript.

## Data and materials availability

All data needed to evaluate the conclusions in the paper are present in the paper and the Supplementary Information. Additional data related to this paper may be requested from the authors.

## Supplementary Information

**Fig. S1.**
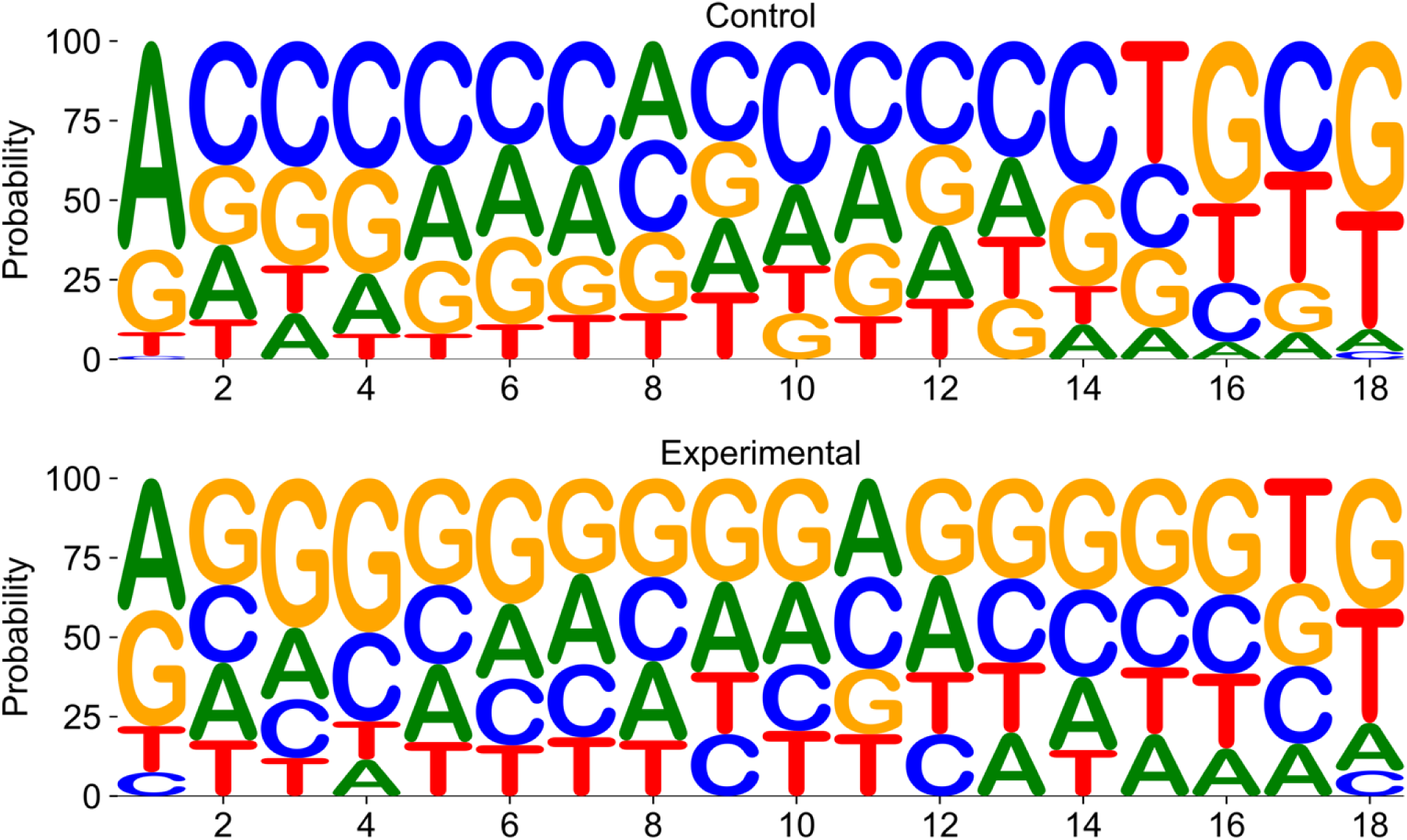
Nucleotide probability at each position across the 18-mer random region in the experimental and control libraries in SELEC round 6. Cytosine is preferred at most positions in the round 6 control (no oxytocin) sequences, whereas guanine is preferred at most positions in the round 6 experimental (with oxytocin) sequences.

**Fig. S2.**
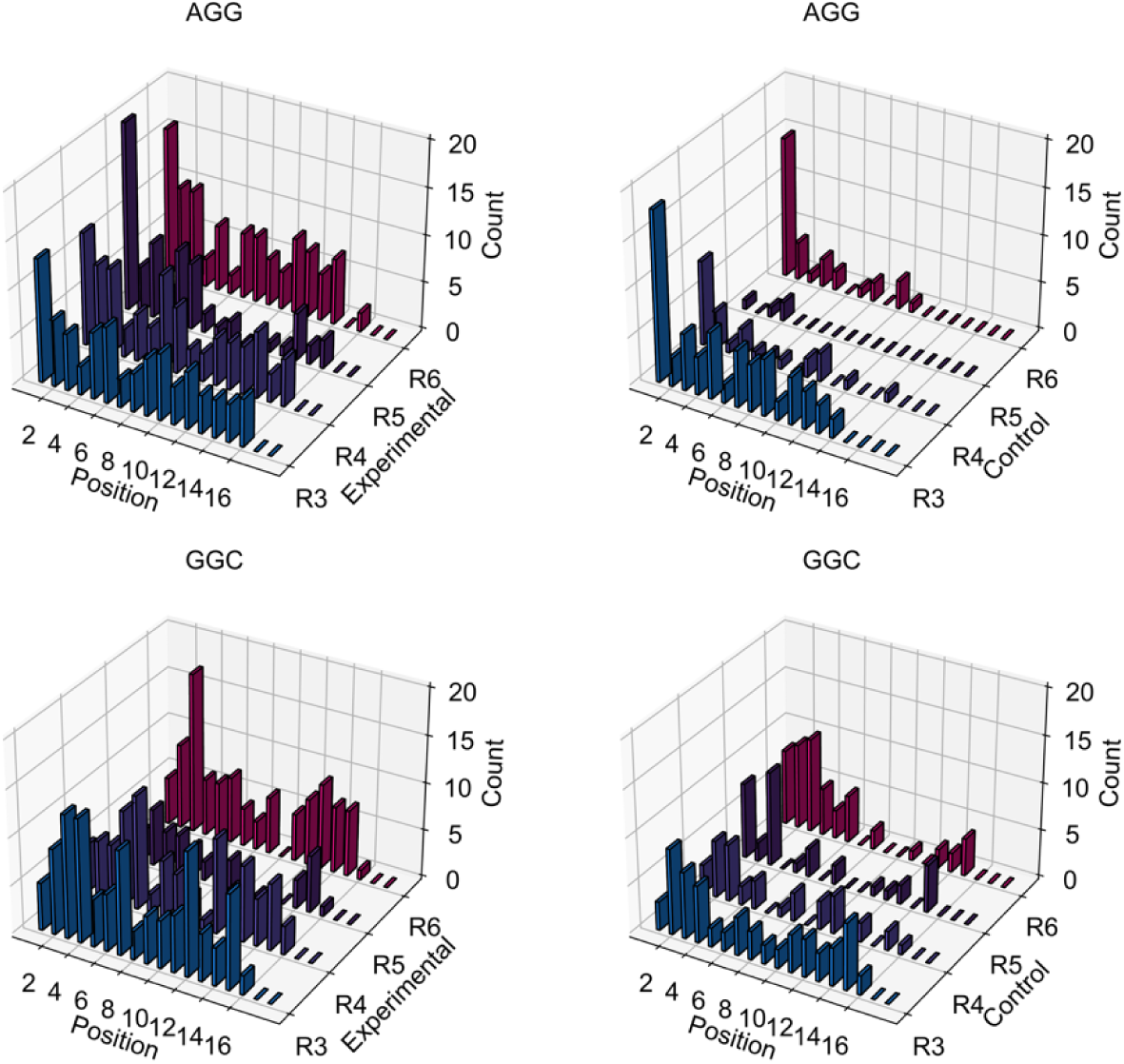
Prevalence of trimers most enriched in the SELEC experimental libraries. AGG and GGC are enriched in experimental libraries over control libraries. Further, these trimers demonstrate positional dependence for the 5’ end of the random recognition region.

**Fig. S3.**
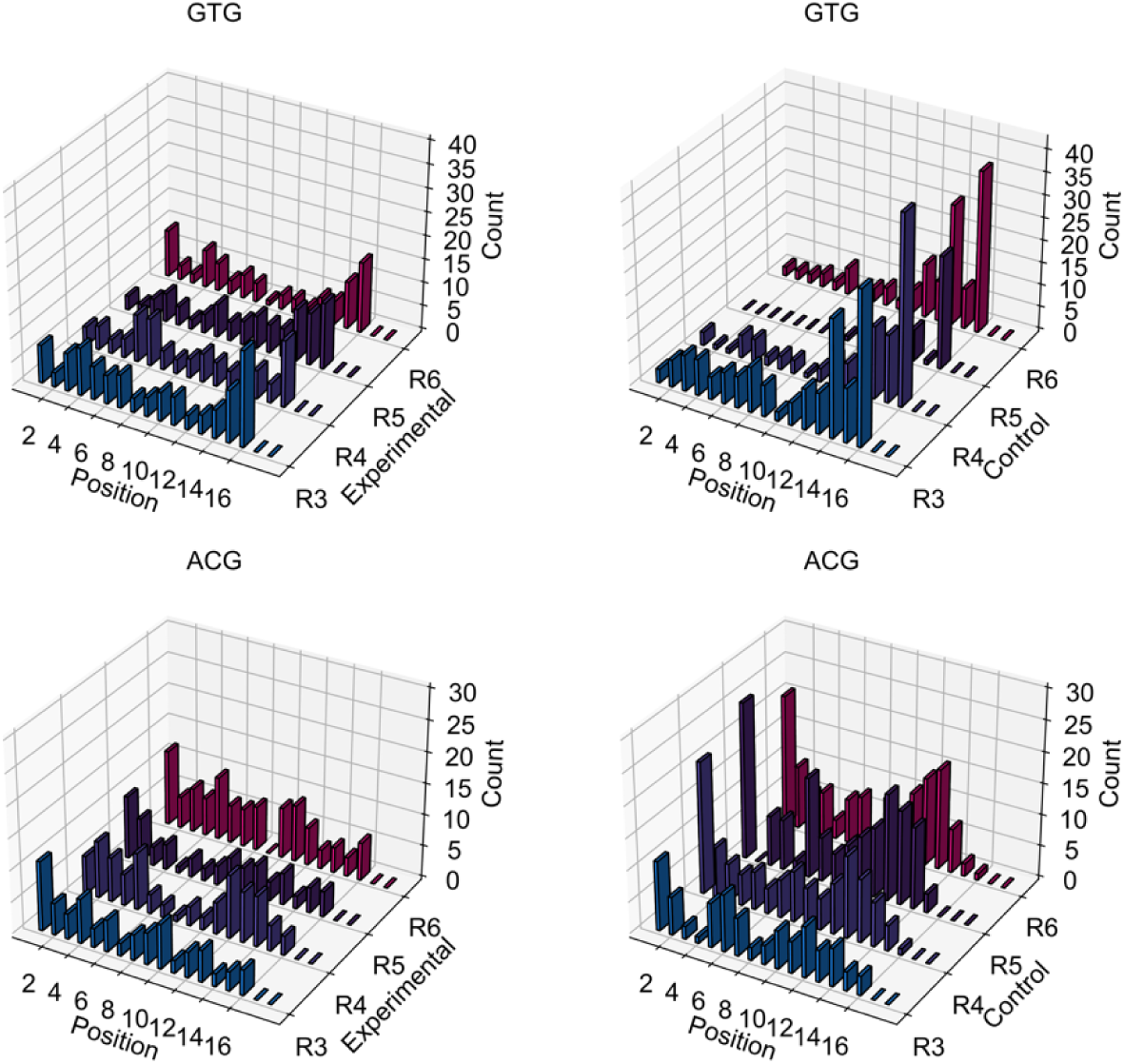
Prevalence of top 10 trimers abundant in SELEC experimental and control libraries. GTG is enriched in control libraries over experimental libraries at position 14 of the 18-mer random region. ACG is enriched in control libraries over experimental libraries at position 1 of the 18-mer random region.

**Fig S4.**
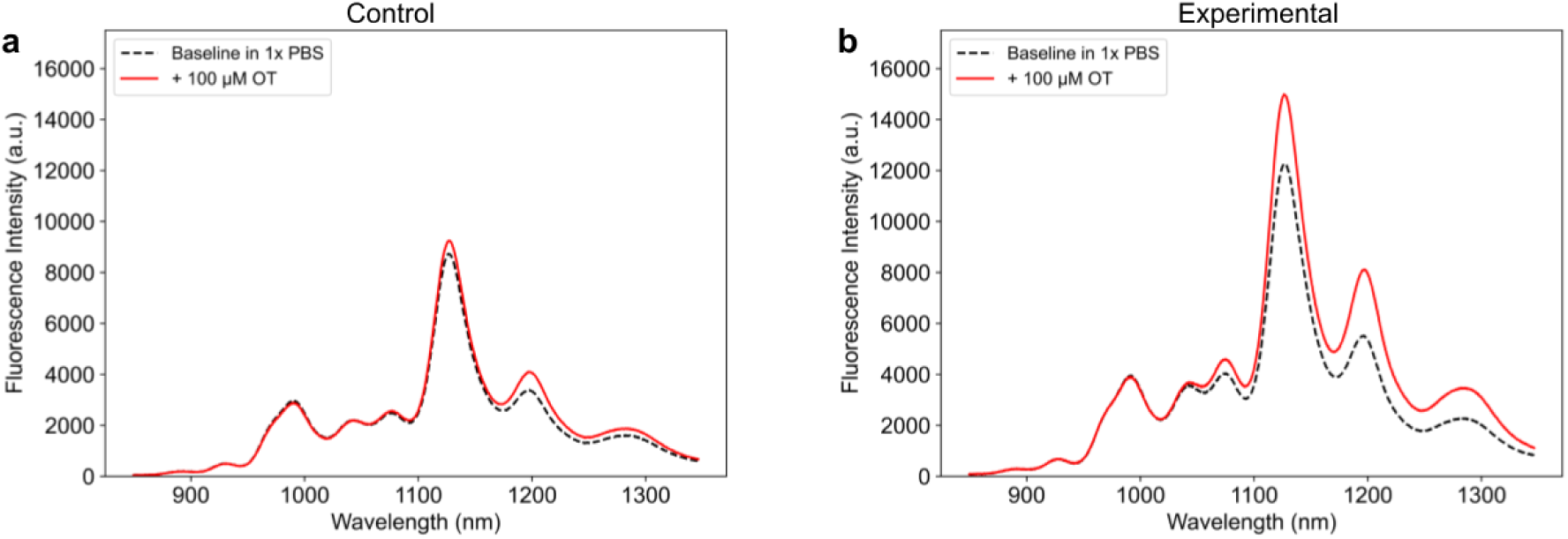
Fluorescence spectra of evolved ssDNA-SWCNT constructs from SELEC experimental and control libraries. (a) C6#4, ssDNA-SWCNT and (b) E6#7 ssDNA-SWCNT before (black, dotted) and after (red) addition of 100 μM oxytocin.

**Fig S5.**
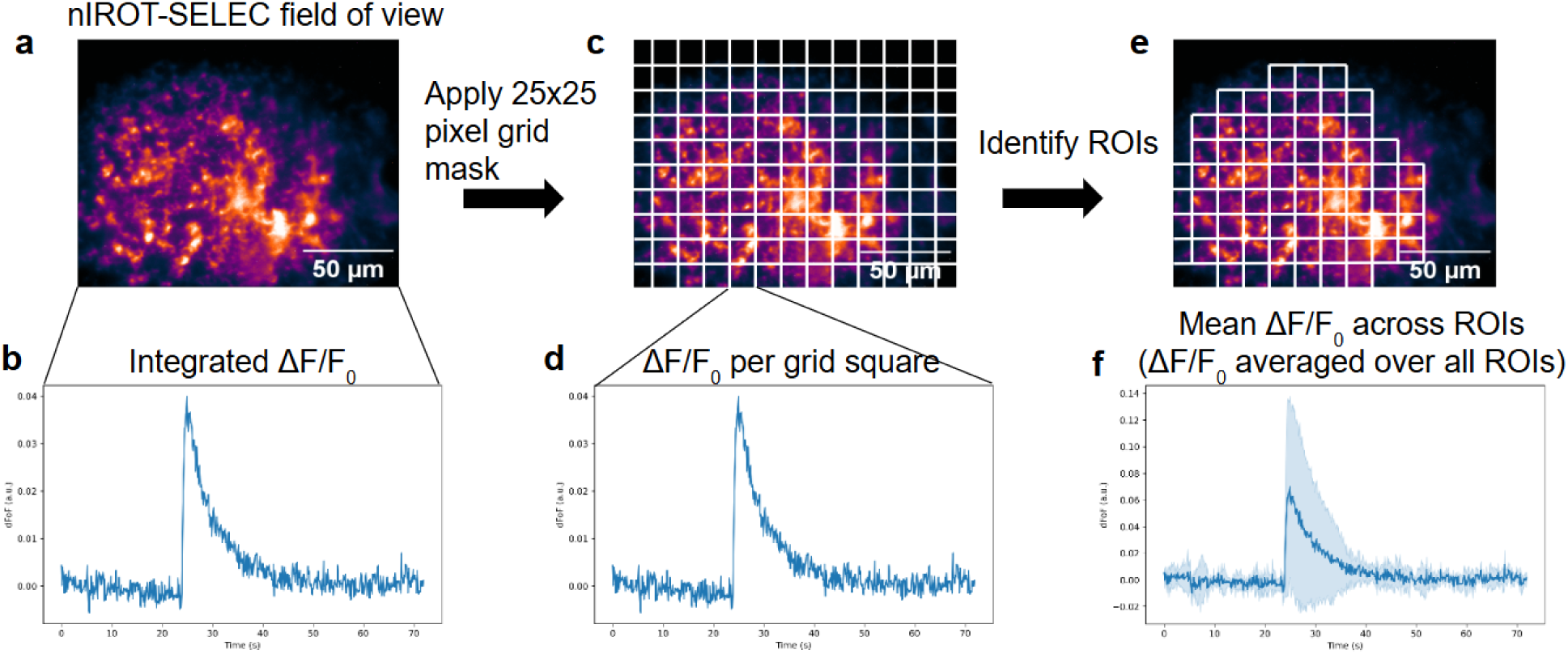
Image processing and data analysis of nanosensor fluorescence response. (a) Imaging movie files to be processed showing the entire field of view (175 µm by 140 µm). (b) Integrated ΔF/F_0_ (ΔF/F_0_ for the entire field of view) was calculated as ΔF/F_0_ = (F-F_0_) /F_0_, where F_0_ is the average intensity for the first 5% of frames and F is the dynamic fluorescence intensity. This will be further averaged over three stimulation replicates. (c) A 25×25 pixel (corresponding to 6.8 µm by 6.8 µm) grid mask was applied to the image stack. (d) ΔF/F_0_ was calculated for each grid square. (e) Grid squares were identified as ROIs if the F around time of stimulation (200 frames) is 3 standard deviations above the baseline F_0_ activity. Note that grid masks here are only for schematic purposes and do not represent actual grids identified by the analysis. (f) ΔF/F_0_ was averaged over all identified ROIs, corresponding to the mean ΔF/F_0_ across ROIs. The number of ROIs and the mean ΔF/F_0_ across ROIs are averaged over at least three experimental replicates. Solid line represents mean and shades represent standard deviations.

**Fig S6.**
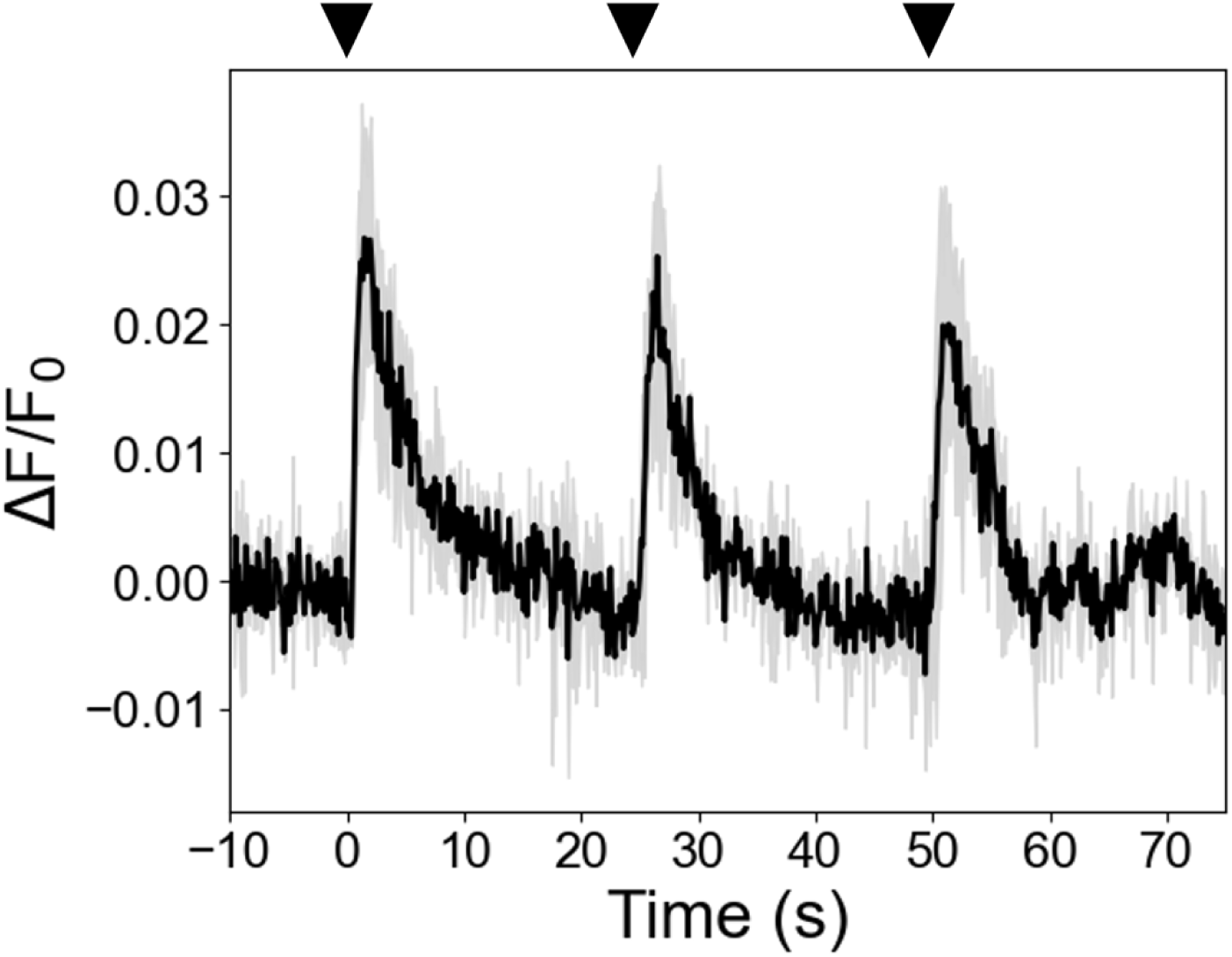
Integrated ΔF/F_0_ of nIROT-SELEC labeled brain slices upon repeated electrical stimulation. 0.5 mA stimulation was applied to a slice containing the paraventricular nucleus of the hypothalamus. Stimulation was repeated every 25 seconds. Nanosensor integrated ΔF/F_0_ was 0.023 ± 0.008, 0.020 ± 0.006 and 0.019 ± 0.007 (means ± SD) for the 1st, 2nd, and 3rd stimulation, respectively, over 2 stimulation replicates. The solid black line represents the mean and the gray shading represents the standard deviation.

**Fig S7.**
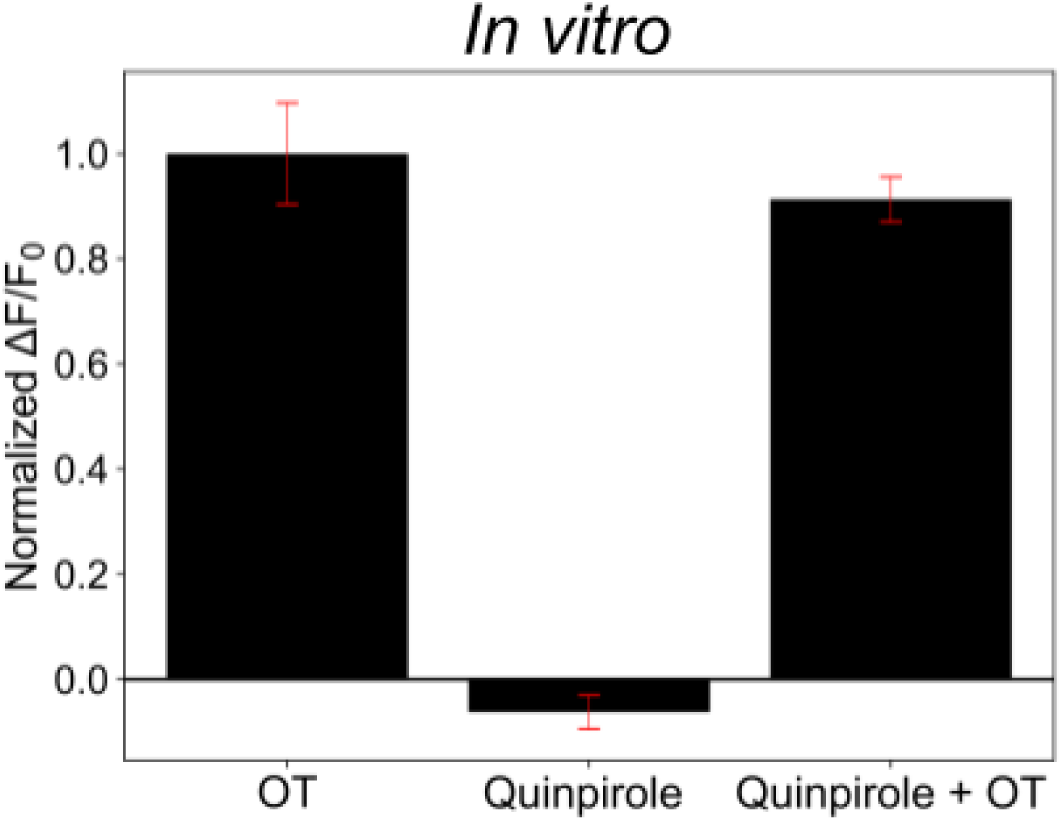
Normalized ΔF/F_0_ of nIROT-SELEC in response to oxytocin, quinpirole, and quinpirole + oxytocin *in vitro*. The effect of 2 µM quinpirole application on nIROT-SELEC was investigated *in vitro*. ΔF/F_0_ was measured in response to oxytocin with no prior quinpirole incubation, quinpirole alone, and oxytocin with prior quinpirole incubation. Before quinpirole application, nIROT-SELEC normalized response to oxytocin was 1.00 ± 0.10. Following quinpirole application, nIROT-SELEC normalized response to oxytocin was 0.91 ± 0.04. Measurements were averaged over 3 technical replicates. Error bars denote standard deviation.

**Fig S8.**
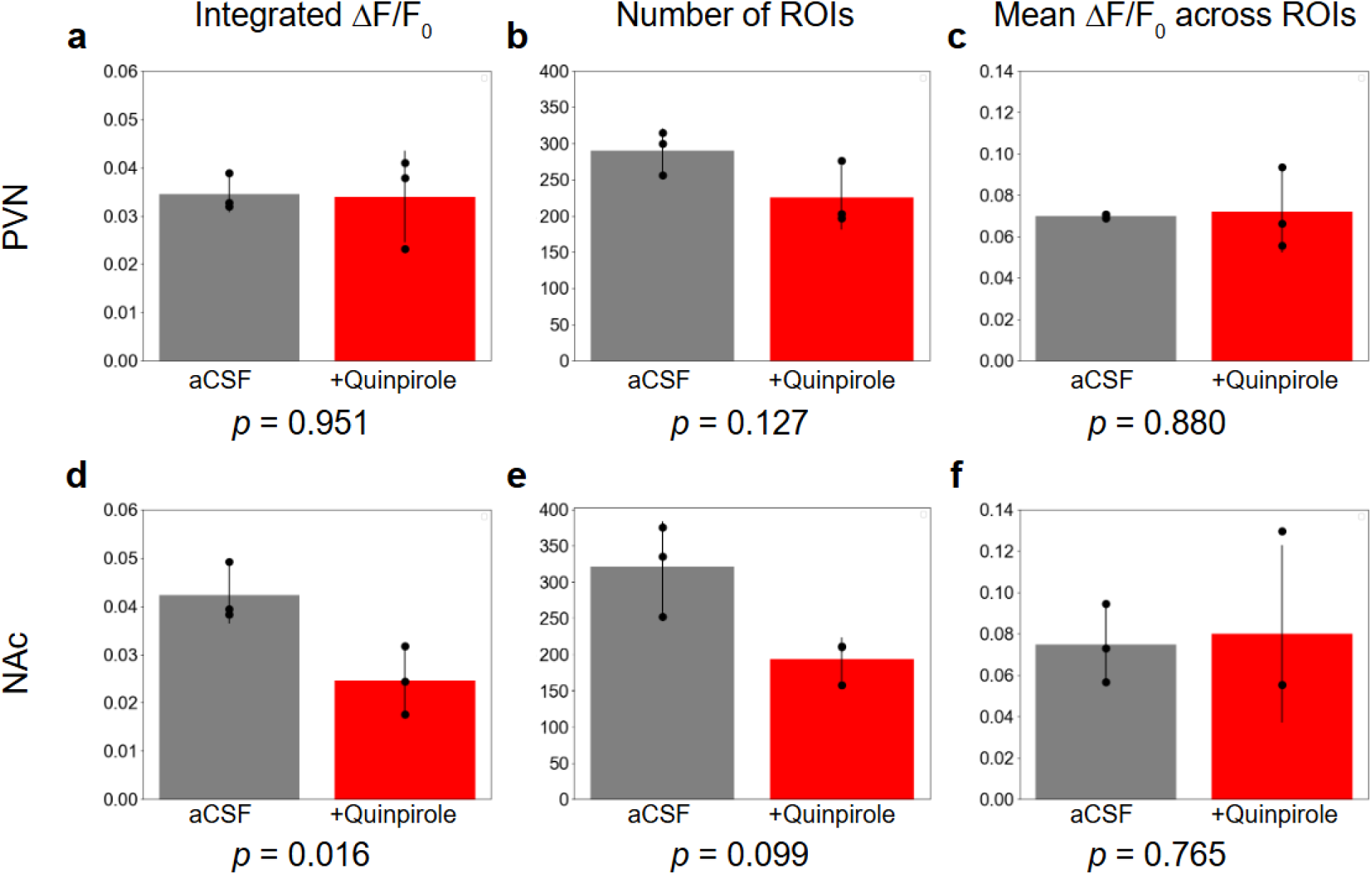
Effect of quinpirole application for oxytocin imaging in acute brain slices of mice. (a) Integrated ΔF/F_0_, (b) number of ROIs, and (c) mean ΔF/F_0_ across ROIs in the paraventricular nucleus (PVN) of the hypothalamus. (d) Integrated ΔF/F_0_, (e) mean number of ROIs, and (f) mean ΔF/F_0_ across ROIs in the nucleus accumbens (NAc). For all figures, each point corresponds to the averaged ΔF/F_0_ or number of ROIs values over three stimulation replicates. *p* values were obtained by performing paired t-tests on values before and after quinpirole applications. All bar graphs show the mean with error bars denoting the standard deviation. n = 3 animals.

**Fig S9.**
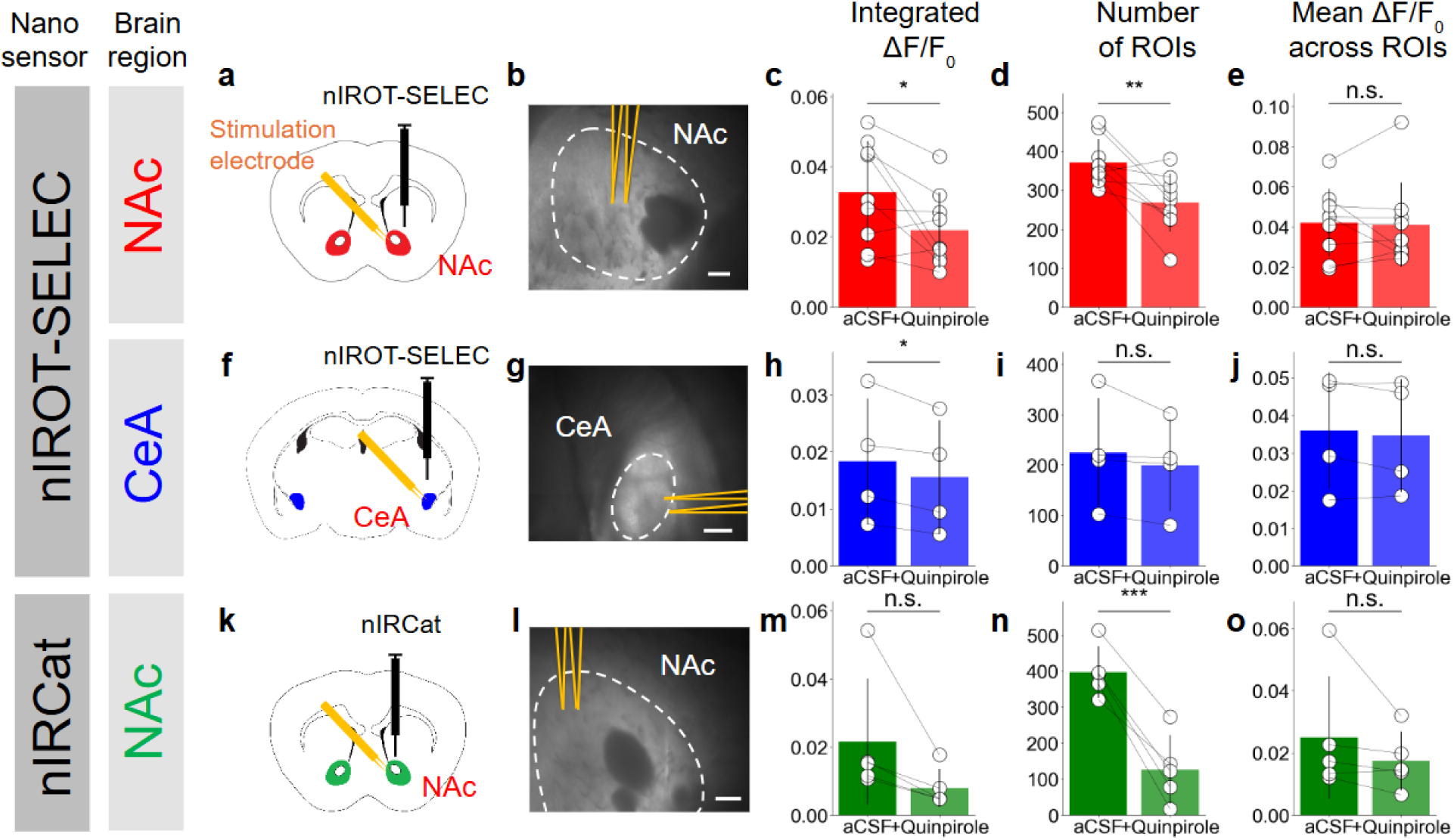
nIROT-SELEC detects neuromodulators *ex vivo* in prairie voles. **(a-e)** nIROT-SELEC was applied to the nucleus accumbens (NAc), **(f-j)** to the central nucleus of the amygdala (CeA), and **(k-o)** nIRCat was applied to the nucleus accumbens. **(a, f, k)** Schematic of brain regions imaged with bipolar stimulator (yellow). **(b, g, l)** Bright field images of brain regions (surrounded by white dash) imaged with bipolar stimulator (surrounded with yellow). Scale bars represent 100 µm. **(c, h, m)** The maximum value of integrated ΔF/F_0_ in aCSF and in aCSF with 2 µM quinpirole. Error bars represent SD. **(c)** [n = 9; *P* = 0.0354] **(h)** [n = 4; *P* = 0.0335] **(m)** [n = 5; *P* = 0.0625] **(d, i, n)** The number of ROIs in aCSF and in aCSF with quinpirole. **(d)** [n = 9; *P* = 0.0082]. **(i)** [n = 4; *P* = 0.1635] **(n)** [n = 5; *P* = 0.0008] **(e, j, o)** Mean ΔF/F_0_ across ROIs in aCSF and in aCSF with quinpirole. Error bars represent SD. **(e)** [n = 9; *P* = 0.8203]. **(j)** [n = 4; *P* = 0.3515] **(o)** [n = 5; *P* = 0.1250]. **(c-e, h-j, m-o)** *p* values were obtained by performing paired t-tests on values before and after quinpirole applications. Statistical significance annotation follows *** for *p* < 0.001, ** for *p* < 0.01, * for *p* < 0.05, and n.s. for p > 0.05.

**Table S1.** Ten most frequent ssDNA trimers from rounds 3 to 6 in experimental SELEC. Trimers are denoted in the order of descending frequency.

| Experimental<br>Trimers | **Roun**<br>**d 3** | **Round**<br>**4** | **Round**<br>**5** | **Round**<br>**6** | **Total** |
| --- | --- | --- | --- | --- | --- |
| **GTG** | 108 | 95 | 90 | 83 | 376 |
| **AGG** | 87 | 87 | 65 | 98 | 337 |
| **GGC** | 107 | 92 | 41 | 93 | 333 |
| **GGT** | 78 | 87 | 82 | 78 | 325 |
| **GGG** | 97 | 90 | 59 | 78 | 324 |
| **TGG** | 96 | 75 | 77 | 75 | 323 |
| **GGA** | 75 | 96 | 64 | 87 | 322 |
| **ACG** | 71 | 83 | 61 | 94 | 309 |
| **CGG** | 87 | 103 | 35 | 80 | 305 |
| **GCA** | 81 | 67 | 46 | 97 | 291 |

**Table S2.** Ten most frequent ssDNA trimers from rounds 3 to 6 in control SELEC. Trimers are denoted in the order of descending frequency.

| Control Trimers | Round 3 | Round 4 | Round 5 | Round 6 | Total |
| --- | --- | --- | --- | --- | --- |
| **CAC** | 33 | 104 | 277 | 109 | 523 |
| **ACA** | 49 | 88 | 228 | 95 | 460 |
| **GTG** | 153 | 117 | 36 | 126 | 432 |
| **ACG** | 75 | 106 | 122 | 124 | 427 |
| **ACC** | 32 | 93 | 156 | 87 | 368 |
| **GAC** | 81 | 84 | 80 | 118 | 363 |
| **CCA** | 30 | 80 | 154 | 93 | 357 |
| **CCG** | 42 | 84 | 117 | 95 | 338 |
| **CGT** | 75 | 83 | 64 | 98 | 320 |
| **GCA** | 40 | 79 | 111 | 87 | 317 |

**Table S3.**
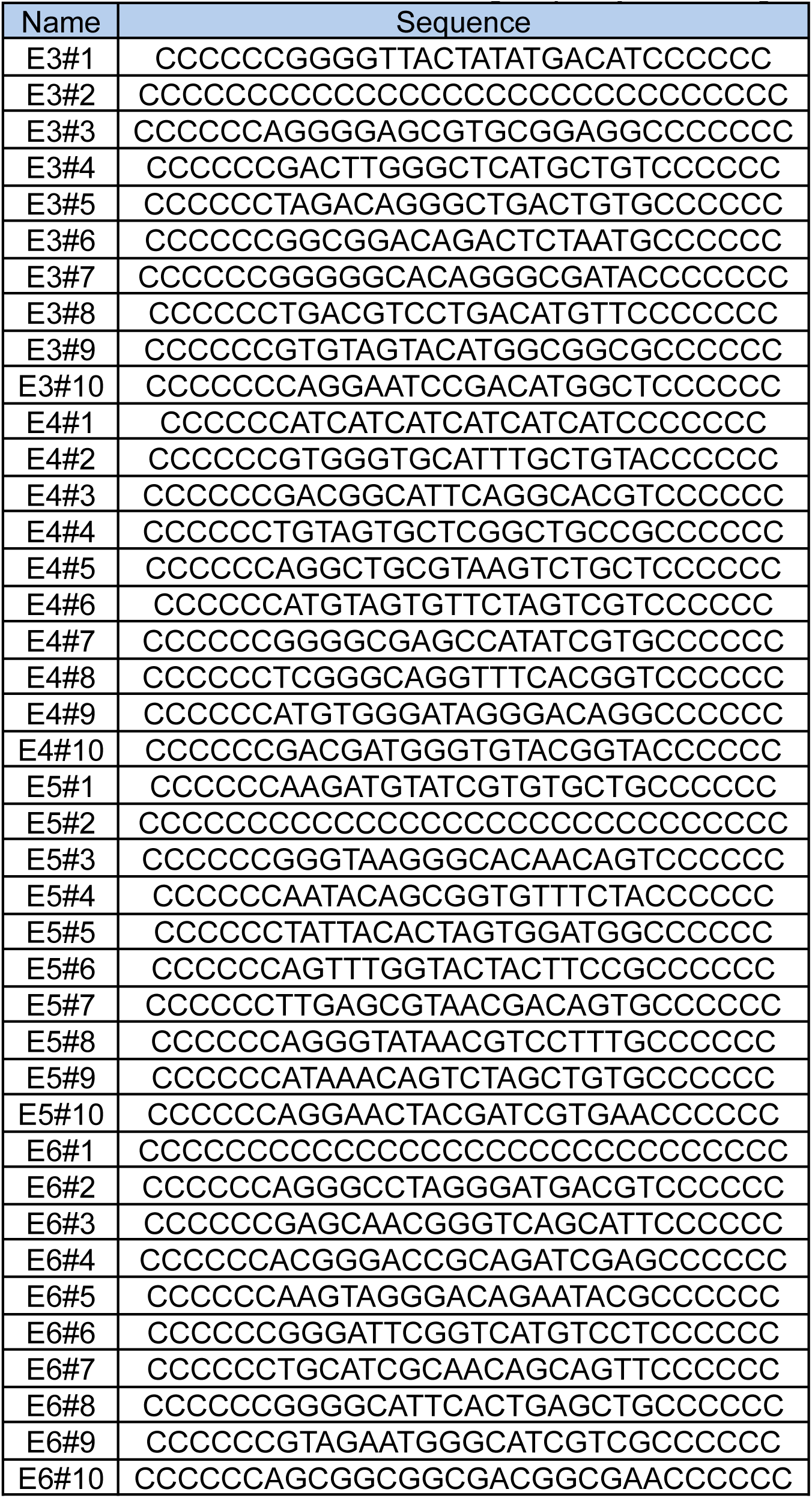
Ten most frequent ssDNA sequences from rounds 3 to 6 in experimental SELEC groups. #1 to #10 are denoted in the order of descending frequency. Primer region is omitted.

## References

1. Froemke, R. C. & Young, L. J. Oxytocin, Neural Plasticity, and Social Behavior. Annu Rev Neurosci 44, 359–381 (2021).

2. Carter, C. S. et al. Is Oxytocin “Nature’s Medicine”? Pharmacol Rev 72, 829 (2020).

3. Jurek, B. & Neumann, I. D. The Oxytocin Receptor: From Intracellular Signaling to Behavior. Physiological Reviews 98, 1805–1908 (2018).

4. Donaldson, Z. R. & Young, L. J. Oxytocin, Vasopressin, and the Neurogenetics of Sociality. Science 322, 900–904 (2008).

5. Winslow, J. T. & Insel, T. R. Neuroendocrine basis of social recognition. Curr Opin Neurobiol 14, 248–253 (2004).

6. Young, L. J. & Wang, Z. The neurobiology of pair bonding. Nat Neurosci 7, 1048–1054 (2004).

7. Yoshida, M. et al. Evidence That Oxytocin Exerts Anxiolytic Effects via Oxytocin Receptor Expressed in Serotonergic Neurons in Mice. J. Neurosci. 29, 2259–2271 (2009).

8. Kirsch, P. Oxytocin in the socioemotional brain: implications for psychiatric disorders. Dialogues in Clinical Neuroscience 17, 463 (2015).

9. Yoon, S. & Kim, Y.-K. Possible oxytocin-related biomarkers in anxiety and mood disorders. Prog Neuropsychopharmacol Biol Psychiatry 116, 110531 (2022).

10. Peled-Avron, L., Abu-Akel, A. & Shamay-Tsoory, S. Exogenous effects of oxytocin in five psychiatric disorders: a systematic review, meta-analyses and a personalized approach through the lens of the social salience hypothesis. Neuroscience & Biobehavioral Reviews 114, 70–95 (2020).

11. Freeman, S. M. & Young, L. J. Comparative Perspectives on Oxytocin and Vasopressin Receptor Research in Rodents and Primates: Translational Implications. Journal of Neuroendocrinology 28, (2016).

12. Anacker, A. & Beery, A. Life in groups: the roles of oxytocin in mammalian sociality. Frontiers in Behavioral Neuroscience 7, (2013).

13. Walum, H. & Young, L. J. The neural mechanisms and circuitry of the pair bond. Nature Reviews Neuroscience 19, 643–654 (2018).

14. Inutsuka, A., Ino, D. & Onaka, T. Detection of neuropeptides in vivo and open questions for current and upcoming fluorescent sensors for neuropeptides. Peptides 136, 170456 (2021).

15. Chefer, V. I., Thompson, A. C., Zapata, A. & Shippenberg, T. S. Overview of Brain Microdialysis. Current Protocols in Neuroscience 47, (2009).

16. Asai, K., Ivandini, T. A. & Einaga, Y. Continuous and selective measurement of oxytocin and vasopressin using boron-doped diamond electrodes. Sci Rep 6, 32429 (2016).

17. Valtcheva, S. et al. Neural circuitry for maternal oxytocin release induced by infant cries. Nature 621, 788–795 (2023).

18. Qian, T. et al. A genetically encoded sensor measures temporal oxytocin release from different neuronal compartments. Nat Biotechnol 41, 944–957 (2023).

19. Jeong, S. et al. High-throughput evolution of near-infrared serotonin nanosensors. Science Advances 5, eaay3771 (2019).

20. Heller, D. A., Baik, S., Eurell, T. E. & Strano, M. S. Single-Walled Carbon Nanotube Spectroscopy in Live Cells: Towards Long-Term Labels and Optical Sensors. Advanced Materials 17, 2793–2799 (2005).

21. Beyene, A. G. et al. Imaging striatal dopamine release using a nongenetically encoded near infrared fluorescent catecholamine nanosensor. Science Advances 5, eaaw3108–eaaw3108 (2019).

22. Bulumulla, C. et al. Visualizing synaptic dopamine efflux with a 2D composite nanofilm. Elife 11, e78773 (2022).

23. Dinarvand, M. et al. Near-Infrared Imaging of Serotonin Release from Cells with Fluorescent Nanosensors. Nano Lett. 19, 6604–6611 (2019).

24. Mun, J. et al. Near-infrared nanosensors enable optical imaging of oxytocin with selectivity over vasopressin in acute mouse brain slices. Proceedings of the National Academy of Sciences 121, e2314795121 (2024).

25. Landry, M. P. et al. Comparative Dynamics and Sequence Dependence of DNA and RNA Binding to Single Walled Carbon Nanotubes. J Phys Chem C Nanomater Interfaces 119, 10048–10058 (2015).

26. Acinas, S. G., Sarma-Rupavtarm, R., Klepac-Ceraj, V. & Polz, M. F. PCR-Induced Sequence Artifacts and Bias: Insights from Comparison of Two 16S rRNA Clone Libraries Constructed from the Same Sample. Appl Environ Microbiol 71, 8966–8969 (2005).

27. Tu, X., Manohar, S., Jagota, A. & Zheng, M. DNA sequence motifs for structure-specific recognition and separation of carbon nanotubes. Nature 460, 250–253 (2009).

28. Nanot, S. et al. Single-Walled Carbon Nanotubes. in Springer Handbook of Nanomaterials 105–146 (Springer, Berlin, Heidelberg, 2013). doi:10.1007/978-3-642-20595-8_4.

29. Pinals, R. L. et al. Rapid SARS-CoV-2 Spike Protein Detection by Carbon Nanotube-Based Near-Infrared Nanosensors. Nano Lett. 21, 2272–2280 (2021).

30. Piekarski, D. J., Boivin, J. R. & Wilbrecht, L. Ovarian Hormones Organize the Maturation of Inhibitory Neurotransmission in the Frontal Cortex at Puberty Onset in Female Mice. Curr Biol 27, 1735–1745.e3 (2017).

31. Yang, S. J., Bonis-O’Donnell, J. T. D., Beyene, A. G. & Landry, M. P. Near-infrared catecholamine nanosensors for high spatiotemporal dopamine imaging. Nat Protoc 16, 3026–3048 (2021).

32. Godin, A. G. et al. Single-nanotube tracking reveals the nanoscale organization of the extracellular space in the live brain. Nat Nanotechnol 12, 238–243 (2017).

33. Young, L. Oxytocin: Synthesis, Secretion and Reproductive Functions. Knobil and Neill, Physiology of Reproduction. in (2006).

34. Yang, S. J. et al. Synaptic scale dopamine disruption in Huntington’s Disease model mice imaged with near infrared catecholamine nanosensors. 2022.09.19.508617 Preprint at 10.1101/2022.09.19.508617 (2022).

35. Melis, M. R., Succu, S., Mascia, M. S., Cortis, L. & Argiolas, A. Extra-cellular dopamine increases in the paraventricular nucleus of male rats during sexual activity. European Journal of Neuroscience 17, 1266–1272 (2003).

36. Kendrick, K. M. Microdialysis measurement of in vivo neuropeptide release. Journal of Neuroscience Methods 34, 35–46 (1990).

37. Young, L. J., Lim, M. M., Gingrich, B. & Insel, T. R. Cellular Mechanisms of Social Attachment. Hormones and Behavior 40, 133–138 (2001).

38. Dölen, G., Darvishzadeh, A., Huang, K. W. & Malenka, R. C. Social reward requires coordinated activity of nucleus accumbens oxytocin and serotonin. Nature 501, 179–184 (2013).

39. Parsons, L. H. & Justice Jr., J. B. Extracellular Concentration and In Vivo Recovery of Dopamine in the Nucleus Accumbens Using Microdialysis. Journal of Neurochemistry 58, 212–218 (1992).

40. Pettit, H. O. & Justice, J. B. Dopamine in the nucleus accumbens during cocaine self-administration as studied by in vivo microdialysis. Pharmacology Biochemistry and Behavior 34, 899–904 (1989).

41. Parsons, L. H. & Justice, J. B. Perfusate serotonin increases extracellular dopamine in the nucleus accumbens as measured by in vivo microdialysis. Brain Research 606, 195–199 (1993).

42. Kenkel, W. M., Gustison, M. L. & Beery, A. K. A Neuroscientist’s Guide to the Vole. Current Protocols 1, e175 (2021).

43. Beery, A. K., Christensen, J. D., Lee, N. S. & Blandino, K. L. Specificity in Sociality: Mice and Prairie Voles Exhibit Different Patterns of Peer Affiliation. Front Behav Neurosci 12, 50 (2018).

44. Beery, A. K. & Shambaugh, K. L. Comparative Assessment of Familiarity/Novelty Preferences in Rodents. Front Behav Neurosci 15, 648830 (2021).

45. Schweinfurth, M. K. et al. Do female Norway rats form social bonds? Behavioral Ecology and Sociobiology 71, 98 (2017).

46. Fletcher, G. J. O., Simpson, J. A., Campbell, L. & Overall, N. C. Pair-Bonding, Romantic Love, and Evolution: The Curious Case of Homo sapiens. Perspect Psychol Sci 10, 20–36 (2015).

47. Schacht, R. & Kramer, K. L. Are We Monogamous? A Review of the Evolution of Pair-Bonding in Humans and Its Contemporary Variation Cross-Culturally. Frontiers in Ecology and Evolution 7, (2019).

48. Numan, M. & Young, L. J. Neural mechanisms of mother-infant bonding and pair bonding: Similarities, differences, and broader implications. Horm Behav 77, 98–112 (2016).

49. Labouesse, M. A., Cola, R. B. & Patriarchi, T. GPCR-Based Dopamine Sensors—A Detailed Guide to Inform Sensor Choice for In Vivo Imaging. International Journal of Molecular Sciences 21, (2020).

50. Hall, H. et al. Distribution of D1- and D2-Dopamine Receptors, and Dopamine and Its Metabolites in the Human Brain. Neuropsychopharmacology 11, 245–256 (1994).

51. Lukas, D. & Clutton-Brock, T. H. The Evolution of Social Monogamy in Mammals. Science 341, 526–530 (2013).

52. Donaldson, Z. R. We’re the Same… but Different: Addressing Academic Divides in the Study of Brain and Behavior. Frontiers in Behavioral Neuroscience 4, (2010).

53. Phelps, S. M., Campbell, P., Zheng, D.-J. & Ophir, A. G. Beating the boojum: Comparative approaches to the neurobiology of social behavior. Neuropharmacology 58, 17–28 (2010).

54. Taborsky, M. et al. Taxon matters: promoting integrative studies of social behavior: NESCent Working Group on Integrative Models of Vertebrate Sociality: Evolution, Mechanisms, and Emergent Properties. Trends in Neurosciences 38, 189–191 (2015).

55. Liu, Y. & Wang, Z. X. Nucleus accumbens oxytocin and dopamine interact to regulate pair bond formation in female prairie voles. Neuroscience 121, 537–544 (2003).

56. Xiao, L., Priest, M. F., Nasenbeny, J., Lu, T. & Kozorovitskiy, Y. Biased Oxytocinergic Modulation of Midbrain Dopamine Systems. Neuron 95, 368–384.e5 (2017).

57. Hung, L. W. et al. Gating of social reward by oxytocin in the ventral tegmental area. Science 357, 1406–1411 (2017).

58. Klinger, M. E., Miller, R. A., Wilbrecht, L. & Landry, M. P. Optical Fibers Functionalized with Single-Walled Carbon Nanotubes for Flexible Fluorescent Catecholamine Detection. bioRxiv 2024.07.09.602792 (2024) doi:10.1101/2024.07.09.602792.

59. Bonis-O’Donnell, J. T. D. et al. Dual Near-Infrared Two-Photon Microscopy for Deep-Tissue Dopamine Nanosensor Imaging. Adv. Funct. Mater. 27, 1702112 (2017).

60. Safaee, M. M. et al. Dual Infrared 2-Photon Microscopy Achieves Minimal Background Deep Tissue Imaging in Brain and Plant Tissues. Advanced Functional Materials n/a, 2404709 (2024).

61. Vaidyanathan, R. & Hammock, E. A. D. Oxytocin receptor dynamics in the brain across development and species. Developmental Neurobiology 77, 143–157 (2017).

62. Beyene, A. G., Demirer, G. S. & Landry, M. P. Nanoparticle-Templated Molecular Recognition Platforms for Detection of Biological Analytes. Curr Protoc Chem Biol 8, 197–223 (2016).

63. Edelstein, A. D. et al. Advanced methods of microscope control using μManager software. J Biol Methods 1, e10 (2014).

